# Ca^2+^-sensor ALG-2 engages ESCRTs to enhance lysosomal membrane resilience to osmotic stress

**DOI:** 10.1101/2024.02.04.578682

**Authors:** Wei Chen, Madeline M. Motsinger, Jiaqian Li, Kevin P. Bohannon, Phyllis I. Hanson

## Abstract

Lysosomes are central players in cellular catabolism, signaling, and metabolic regulation. Cellular and environmental stresses that damage lysosomal membranes can compromise their function and release toxic content into the cytoplasm. Here, we examine how cells respond to osmotic stress within lysosomes. Using sensitive assays of lysosomal leakage and rupture, we examine acute effects of the osmotic disruptant glycyl-L-phenylalanine 2-naphthylamide (GPN). Our findings reveal that widely used concentrations of GPN rupture a small fraction of lysosomes, but surprisingly trigger Ca^2+^ release from nearly all. Chelating cytoplasmic Ca^2+^ using BAPTA makes lysosomes more sensitive to GPN-induced rupture, suggesting that Ca^2+^ plays a role in protecting or rapidly repairing lysosomal membranes. GPN-elicited Ca^2+^ release causes the Ca^2+^-sensor Apoptosis Linked Gene-2 (ALG-2) and ESCRT proteins it interacts with to redistribute onto lysosomes. ALG-2, but not its ESCRT binding-disabled ΔGF^122^ splice variant, improves lysosomal resistance to rupture by GPN or the lysosomotropic drug chlorpromazine. Importantly, directly elevating juxta-lysosomal Ca^2+^ without membrane damage by activating TRPML1 also recruits ALG-2 and ESCRTs, protecting lysosomes from subsequent osmotic rupture. These findings reveal that Ca^2+^, through ALG-2, helps bring ESCRTs to lysosomes to enhance their resilience and maintain organelle integrity in the face of osmotic stress.

**Significance:** As the degradative hub of the cell, lysosomes are full of toxic content that can spill into the cytoplasm. There has been much recent interest in how cells sense and repair lysosomal membrane damage using ESCRTs and cholesterol to rapidly fix “nanoscale damage”. Here, we extend understanding of how ESCRTs contribute by uncovering a preventative role of the ESCRT machinery. We show that ESCRTs, when recruited by the Ca^2+^-sensor ALG-2, play a critical role in stabilizing the lysosomal membrane against osmotically-induced rupture. This finding suggests that cells have mechanisms not just for repairing but also for actively protecting lysosomes from stress-induced membrane damage.

## Introduction

Lysosomes serve as the cell’s primary degradative organelle and play a pivotal role in regulating metabolism and signaling. As a consequence, they face numerous stressors that can damage their membranes, such as osmotic changes, oxidative damage, particulate material and pathogens (1–4). Maintaining lysosomal integrity is crucial to preventing harmful content from leaking into the cytoplasm, and is vital for cell homeostasis and survival. Molecular pathways that repair lysosomal damage have emerged as the focus of great interest in recent years (5–8) but much remains to be learned about the underlying mechanisms and how different responses are coordinated.

Lysosomes regularly encounter osmotic stress, resulting from changes in metabolic activity, accumulation of lysosomotropic compounds, and environmental shifts. Osmotic pressure increases membrane tension, rupturing the lysosome when the tension exceeds a certain threshold. Factors such as lipid peroxidation, abnormal lipid composition and various lysosomal storage disorders can increase susceptibility to damage.

The sensitivity of lysosomes to osmotic stress has been known for decades, initially discovered through *in vitro* studies of lysosomal rupture by membrane-permeant amino acid esters (9–11). Glycyl-L-phenylalanine 2-naphthylamide (GPN) later emerged as an acute and selective lysosomal disruptant because of its metabolism by cathepsin C and is used to permeabilize lysosomes both *in vitro* and in cells (2, 12, 13). Following cleavage by cathepsin C, glycyl-phenylalanine and β-naphthylamide rapidly accumulate to generate osmotic pressure that eventually ruptures the lysosomal membrane (2, 12).

GPN has been extensively employed to deplete lysosomal Ca^2+^ stores (13–18). While there is growing interest in the cross-talk among intracellular Ca^2+^ stores, including between the lysosome and endoplasmic reticulum (ER) after addition of GPN, there remains consensus that GPN induces lysosomal Ca^2+^ release (19, 20). Given the known involvement of Ca^2+^ in initiating membrane repair, this raises questions about whether and how GPN-induced Ca^2+^ release influences cellular responses to lysosomal membrane stress and damage.

Previous studies have shown that the Endosomal Sorting Complex Required for Transport (ESCRT) machinery is recruited to GPN-treated lysosomes, contributing to pH restoration after removal of GPN (5). Similar recruitment has been observed during plasma membrane repair, where ESCRTs accumulate at sites of Ca^2+^ influx through interaction with the penta-EF hand Ca^2+^-binding protein ALG-2 (also known as PDCD6) (21). ALG-2 is a dimer that facilitates assembly of ESCRT and other proteins on membranes in a Ca^2+^-dependent manner in cells (22) and *in vitro* (23). Among ESCRTs, ALG-2 binds to ALIX, TSG101, VPS37B/C, and IST1 (24–30). Functionally, ALG-2 helps repair laser-induced wounds in the plasma membrane (21) and promotes recovery from electroporation (31). It also works together with ESCRTs to maintain the integrity of membranes around expanding *Candida albicans* hyphae as they invade host cells (32). On lysosomes, Ca^2+^-ALG-2 binds the cation-permeant TRPML1 channel and connects it to dynein to promote organellar movement toward the perinuclear region (33, 34). ALG-2 also accumulates with ESCRTs on lysosomes damaged by L-leucyl-L-leucine methyl ester (LLOMe) (5, 23, 35) but is not required for ESCRT recruitment in this setting (35). Whether ALG-2 contributes to lysosomal membrane homeostasis and/or repair is thus not known.

In this study, we characterize cellular responses to the osmotic stress induced by GPN in lysosomes, distinguishing between phases of Ca^2+^ release and subsequent osmotic rupture. We demonstrate that low GPN concentrations trigger Ca^2+^ release without causing leakage or rupture, and that rapid Ca^2+^ chelation increases lysosomal vulnerability to rupture by GPN. Our findings further establish that GPN prompts recruitment of ALG-2 and associated ESCRTs, enhancing lysosomal resilience to osmotic stress. Finally, we show that activating the TRPML1 cation channel recruits ALG-2 and ESCRTs to intact lysosomes, mitigating subsequent damage. These results demonstrate a protective role for lysosomal Ca^2+^ release and ALG-2-ESCRT complexes on lysosomes.

## Results

### Effects of GPN on lysosomal integrity

To explore cellular responses to osmotic stress in lysosomes, we revisited the effects of GPN. Tracking the loss of lysosomal contents can be challenging, especially when leakage is partial or affects only a few lysosomes. We therefore developed a sensitive assay using the membrane impermeant dye SYTOX Green (hereafter referred to as SYTOX) to monitor the collective leakage of a low molecular weight compound from a cell’s lysosomes. After loading this dye into lysosomes using an endocytosis-chase protocol (Fig. 1A&B), we followed its leakage taking advantage of the several hundred-fold amplification in SYTOX fluorescence upon binding to DNA (36). Upon treating cells with GPN at a concentration commonly thought to permeabilize lysosomes (0.2 mM) (5, 13, 37), we noted a gradual increase in nuclear fluorescence as SYTOX escaped and diffused into the nucleus (Fig. 1C, movie 1). By quantifying nuclear fluorescence over time, we established a consistent measure of lysosomal leakage within individual cells (Fig. 1D). Cells not exposed to GPN, but loaded with SYTOX, showed no change in nuclear fluorescence, confirming that imaging and associated phototoxicity did not compromise lysosomal integrity (Fig. 1D&S1). Inhibiting cathepsin C with the cysteine protease inhibitor E64d prevented GPN-induced SYTOX release, supporting the expected requirement for lysosomal proteolysis in osmolyte generation (Fig. 1D&S1). Additionally, we assessed plasma membrane permeability during GPN treatment by adding propidium iodide together with GPN. Nuclear SYTOX increased as expected over 30 min while propidium iodide was not evident until we added 0.1% Triton X-100 (Fig. S2A) confirming that GPN’s effect on SYTOX leakage was specific to lysosomal disruption.

**Figure 1.**
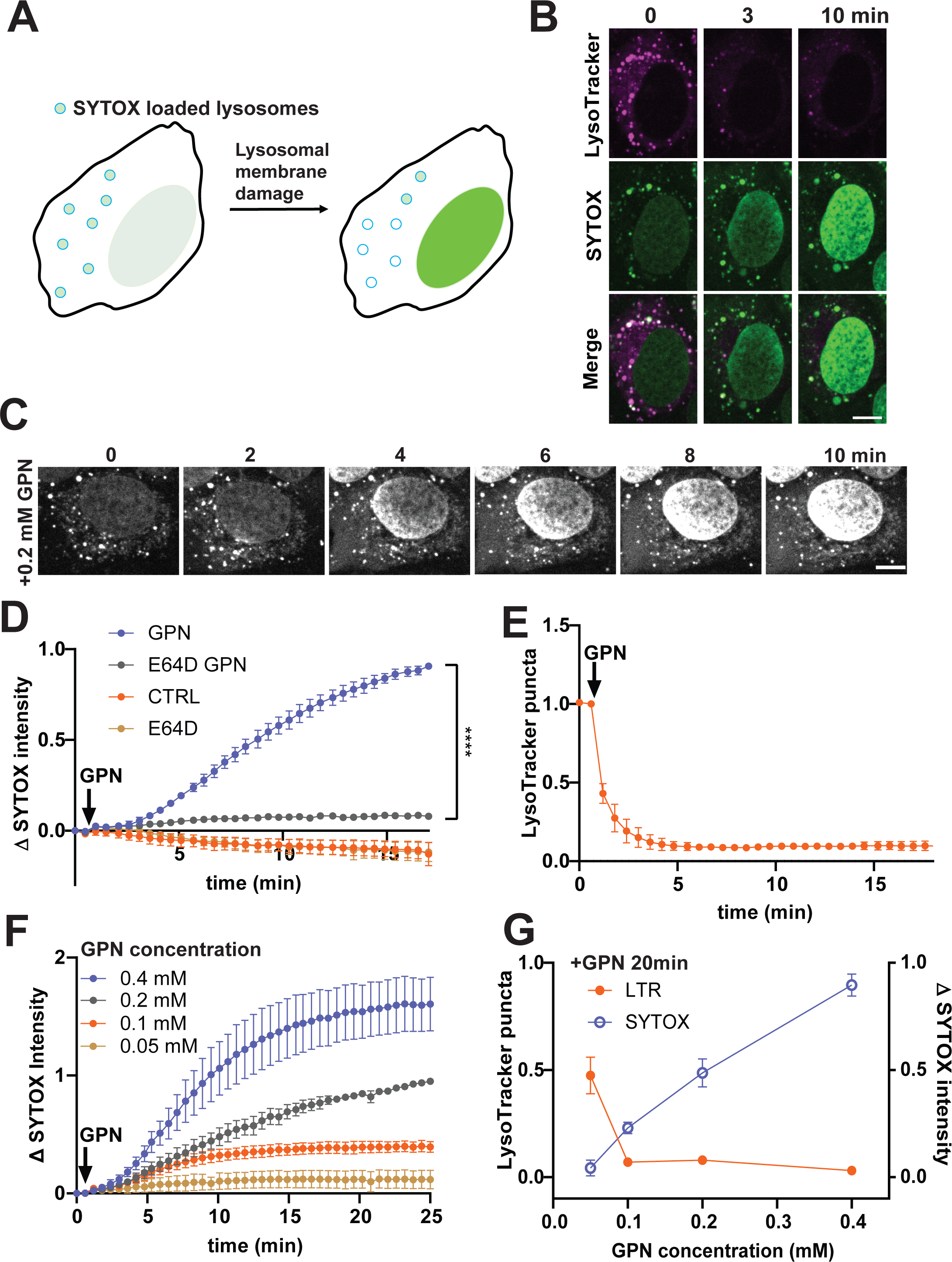
Effects of GPN on lysosomal integrity and pH. (A) Schematic of SYTOX loss assay used to monitor cumulative lysosomal damage in single cells. Lysosomal SYTOX is released upon membrane damage and detected after diffusion into the nucleus. (B) Representative U2OS cell with SYTOX-loaded and Lysotracker-stained lysosomes treated with GPN for indicated time. (C) Representative U2OS cell with SYTOX-loaded lysosomes treated with GPN for indicated time. A whole field view and time course is shown in Movie 1. (D) Effects of 0.2 mM GPN on nuclear SYTOX accumulation in U2OS cells pretreated or not with E64d. Images were captured every 15 s after addition of DMSO or GPN. Changes in nuclear SYTOX intensity (mean ± SEM, normalized to mean nuclear SYTOX intensity change after 20 min of GPN in each experiment; n>100 cells for each condition from 3 independent experiments) were measured for each time point. Kruskal– Wallis test was used to compare changes among different groups at 20 min. **** denotes P<0.0001. (E) U2OS cells stained with Lysotracker Red (LTR) and imaged after addition of 0.2 mM GPN. Y axis, number of LTR puncta (mean ± SEM per field, normalized to the number of LTR puncta at 0 min; more than 100 cells for each condition from 3 independent experiments). (F) Effect of indicated concentrations of GPN on nuclear SYTOX intensity (mean ± SEM, normalized to the mean nuclear SYTOX intensity change in cells treated with 0.2 mM GPN after 25 min in each experiment; n>100 cells for each condition from 3 independent experiments). (G) Comparison of LTR and SYTOX responses to varied concentrations of GPN in U2OS cells. Normalized number of LTR puncta (normalized to the number of LTR puncta at 0 min) and nuclear SYTOX intensity change (mean ± SEM, normalized to the mean nuclear SYTOX intensity change seen using 0.4 mM at 25 min in each experiment; n>100 cells for each condition from 3 independent experiments). Scale bars, 10µm.

Interestingly, despite prolonged exposure to GPN a subset of SYTOX-containing organelles remained unchanged in the cytoplasm (Fig. 1B&C). Concurrent monitoring with Lysotracker Red showed that GPN rapidly increased pH in all lysosomes (Fig. 1B&E), indicating that although all were buffered by GPN some resisted rupture. Given that early *in vitro* studies had established a robust correlation between GPN concentration and liberation of lysosomal enzymes (2), we next assessed the impact of varying GPN doses on lysosomal integrity in cells. Strikingly, doubling the concentration of GPN from 0.2 to 0.4 mM caused significantly more SYTOX release, and further experiments established a dose-dependent effect on lysosomal leakage (Fig. 1F&G, S3). The fact that Lysotracker staining was lost at lower concentrations of GPN than those causing significant SYTOX release is readily explained by considering differences in how these probes leave the lysosome (Fig. 1G). Lysotracker Red is a reversible acid sensitive probe that concentrates in lysosomes but rapidly dissipates when the pH gradient is lost independently of membrane damage, for example following addition of the weak base NH_4_Cl (Fig. S4). SYTOX, in contrast, is membrane impermeant and its loss requires at least transient membrane permeabilization. SYTOX is not released from lysosomes in cells treated with NH_4_Cl (Fig. S4) because different from the metabolites of GPN, NH_4+_ is exported via a specific transporter (38) and does not accumulate to generate osmotic pressure. The observation that substantially higher concentrations of GPN are needed to release SYTOX than to neutralize lysosomal pH raises questions about the extent to which typical GPN concentrations compromise lysosomal integrity.

To further examine lysosomal damage elicited by GPN, we monitored Galectin-3 (GAL3) recruitment (1, 39). Cytoplasmic GAL3 binds β-galactosides present within the lysosomal lumen when they are exposed by membrane damage, leading to accumulation of GAL3 on or within ruptured lysosomes. In cells stably expressing mCherry-GAL3, adding increasing concentrations of GPN revealed a dose-dependent relationship between GPN and the appearance of mCherry-GAL3 puncta (Fig. 2A&B). At typically used GPN concentrations (0.2 mM) the number of GAL3 puncta was small compared to the total number of lysosomes. Paralleling this, loss of 10 kDa dextran was minimal (Fig. S5).

**Figure 2.**
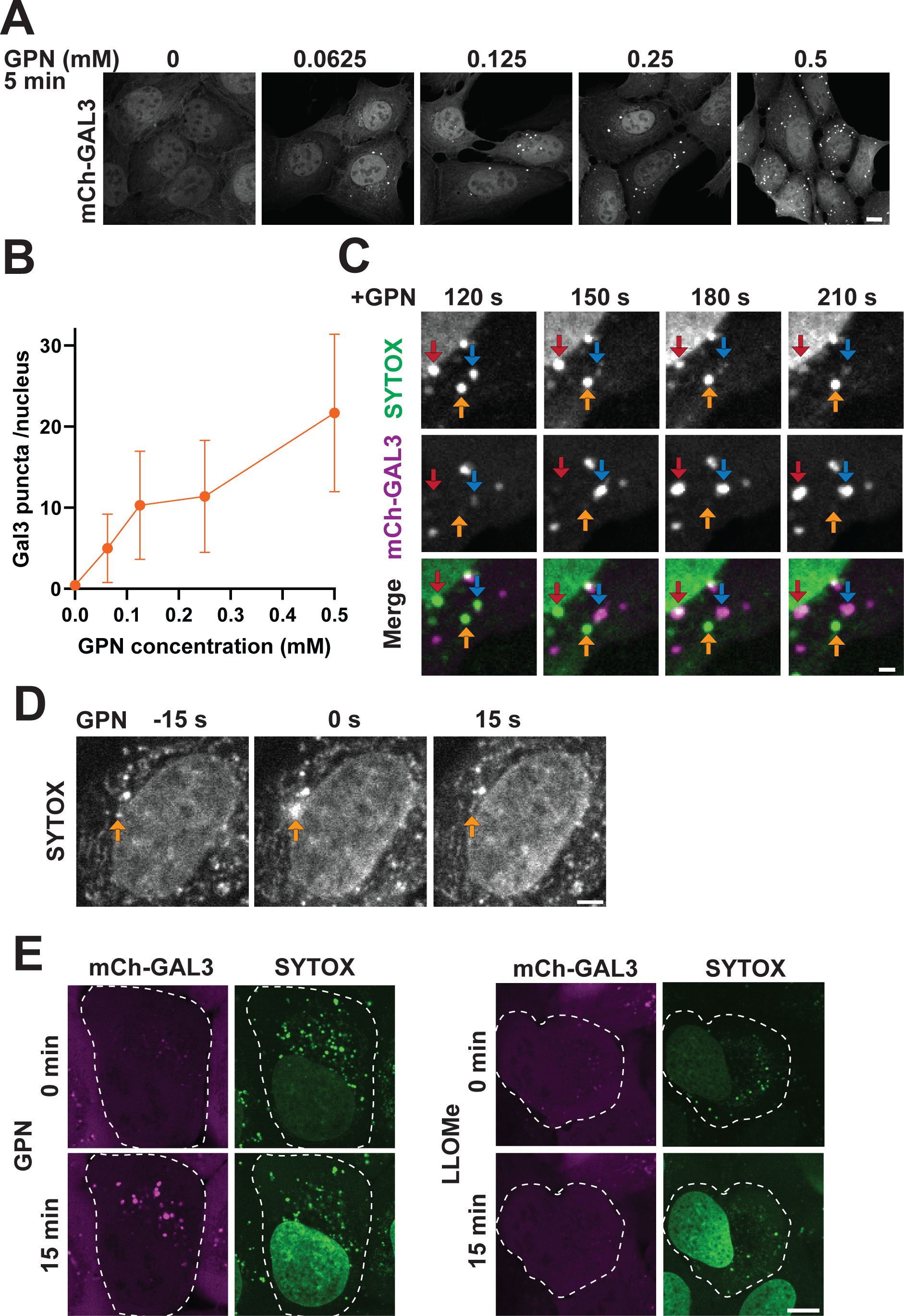
GPN-induced SYTOX loss coincides with recruitment of Galectin-3 to ruptured lysosomes. (A) U2OS cells expressing mCherry-GAL3 were treated with different concentrations of GPN for 5 min and fixed before imaging. (B) Plot of GAL3 puncta per cell (mean ± SEM; n>90 cells for each condition from 3 independent experiments). (C) U2OS cells expressing mCherry-GAL3 and loaded with SYTOX were imaged after addition of 0.2 mM GPN. Different color arrows highlight changes seen on three different lysosomes. Red arrows point to a lysosome that gradually lost SYTOX and gained GAL3. Blue arrows point to a lysosome that lost SYTOX and gained GAL3 abruptly. Yellow arrows point to a lysosome that remained intact. (D) Representative burst of SYTOX release from a single rupturing lysosome. Burst is set at time 0. (E) Representative U2OS cells expressing mCherry-GAL3 and loaded with SYTOX were treated with GPN or LLOME for 15min and imaged live. Scale bars, 10 µm (A), 2 µm (C) and 2.5 µm (D).

To directly compare the effects of GPN on SYTOX loss and mCherry-GAL3 accumulation, we turned to live cell imaging to monitor these events. In cells stably expressing mCherry-GAL3 and incubated with 0.2 mM GPN, we observed simultaneous SYTOX loss and mCherry-GAL3 appearance (Fig. 2C). With faster imaging, we saw occasional bursts of SYTOX release indicative of organelle rupture and content loss (Fig. 2C&D, movie1). These bursts paralleled the increase in SYTOX intensity, suggesting that ruptures underlie the majority of GPN-induced SYTOX release (Fig. S6) although we cannot rule out a small amount of leakage from otherwise intact and GAL3-negative organelles.

Because the abrupt GPN-induced appearance of GAL3 puncta differs from the gradual GAL3 accumulation seen with LLOMe (5), we directly compared effects of these compounds (Fig. 2E). Treatment with GPN led to both SYTOX loss and GAL3 accumulation, while LLOMe induced SYTOX loss without the appearance of GAL3 puncta. Chlorpromazine (CPZ), like GPN, also induced bursts of SYTOX release accompanied by abrupt GAL3 recruitment (Fig. S7A&B), underscoring the varied mechanisms by which lysosomotropic compounds damage lysosomes.

### Ca^2+^ mobilized by GPN is from both lysosomes and the endoplasmic reticulum and helps maintain lysosomal integrity

Given that GPN-induced lysosomal rupture requires a higher concentration of GPN than expected based on earlier studies, we next wondered about its widely exploited effects on lysosomal Ca^2+^. Based on foundational *in vitro* studies of content loss after osmotic rupture (2, 12), it has long been thought that the induced rupture underlies lumenal Ca^2+^ release (13, 15, 17, 19, 20, 40). To explore the relationship between GPN-induced osmotic stress and Ca^2+^ release, we fused a fluorescent Ca^2+^ sensor to the cytoplasmic tail of the lysosomal membrane protein TMEM192 (TMEM192-GCAMP7) to monitor changes in juxta-lysosomal Ca^2+^ (14, 41, 42). We also assessed changes in ER lumenal Ca^2+^ using a low-affinity Ca^2+^ sensor, CEPIA, coupled to an ER retention sequence (43). In experiments with U2OS cells expressing both sensors, adding 0.2 mM GPN rapidly increased TMEM192-GCAMP7 fluorescence. The signal peaked within 1-2 min and did not immediately returning to baseline levels (Fig. 3A), indicating rapid juxta-lysosomal Ca^2+^ mobilization. Concurrently, we observed a decrease in ER lumenal Ca^2+^, confirming significant Ca^2+^ loss from the ER (Fig. 3A).

**Figure 3.**
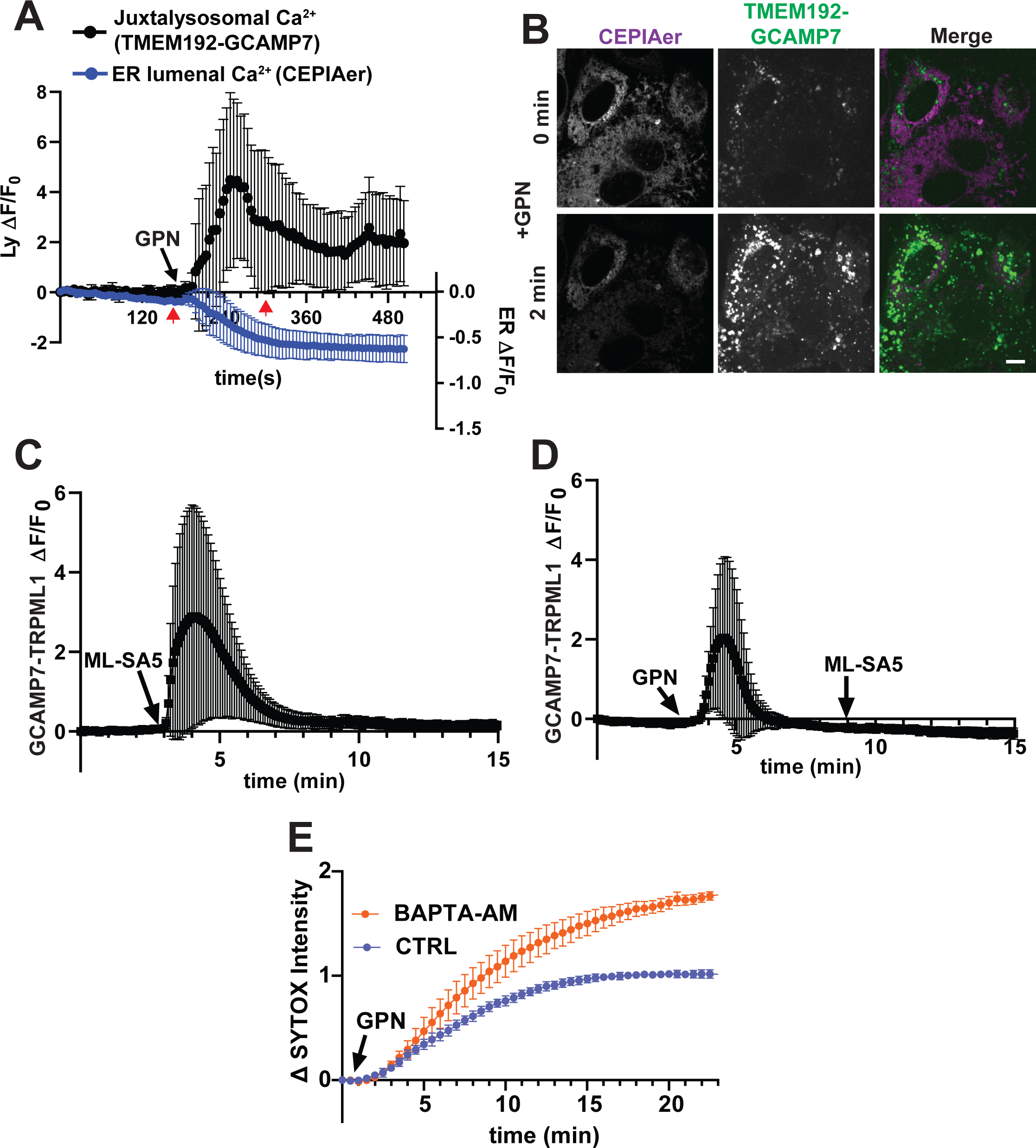
GPN-induced Ca^2+^ signaling and its impact on lysosomal resilience. (A) U2OS cells expressing CEPIAer and TMEM192-GCAMP7 were imaged at 5 s intervals before and after addition of GPN. Intensity changes (ι1F) of CEPIAer and TMEM192-GCAMP7 were measured and normalized to starting value (F0) for each time point. Mean± SD were plotted (n= 13 cells). Red arrows indicate time points captured in representative cells shown in (B). (C) U2OS cells expressing GCAMP7-TRPML1 were imaged in Ca^2+^-free (0 Ca^2+^) imaging solution. ML-SA5 was added at indicated time points without (C) or with (D) GPN pretreatment. Normalized intensity change is plotted (Mean± SD, n=12 and 22 cells). (E) U2OS cells loaded with SYTOX and preincubated with DMSO or BAPTA-AM were imaged at 30 s intervals following GPN treatment. Y axis, change in nuclear SYTOX intensity (mean ± SEM, normalized to the mean value of nuclear SYTOX intensity change in CTRL group at 20 min in each experiment; n= 88 and 131 cells from 3 independent experiments). Unpaired t test was performed for each time point with P<0.05 starting from 11 min.

Interestingly, GPN-induced juxta-lysosomal Ca^2+^ signals were observed across the cell on many more lysosomes than are ruptured by GPN (Fig. 3B). Given recent questions about the specific contributions of lysosomal versus ER Ca^2+^ stores to GPN-elicited Ca^2+^ signals (40), we further explored the effects of GPN on releasable lysosomal Ca^2+^. To do this, we took advantage of a TRPML1 synthetic agonist, ML-SA5 (44), that rapidly and completely releases Ca^2+^ from lysosomes (15). In U2OS cells expressing GCAMP7 fused to TRPML1, ML-SA5 triggered the expected robust Ca^2+^ spike (Fig. 3C). GPN similarly elicited a transient increase in GCAMP7 fluorescence, but there was no further response with subsequent addition of ML-SA5 indicating that GPN had depleted the releasable Ca^2+^ (Fig. 3D). This is consistent with previous reports (15, 20) and establishes that Ca^2+^ escapes lysosomes differently from SYTOX at the typically used concentration of 0.2 mM GPN. Whether Ca^2+^ leaves through transient nanoscale damage or instead a stress-sensitive channel or transporter remains to be determined.

Given the well-established role of Ca^2+^ in membrane repair, we next wondered about the importance of juxta-lysosomal Ca^2+^ in responses to GPN induced stress. In U2OS cells treated with BAPTA-AM, a fast-acting Ca^2+^ chelator, we found that GPN treatment resulted in increased lysosomal SYTOX release (Fig. 3E). This suggests that Ca^2+^ signals help to limit lysosomal leakage, indicating that juxta-lysosomal Ca^2+^ may play a protective role in reducing lysosomal vulnerability to osmotic rupture.

### ALG-2 is required for GPN-triggered ESCRT recruitment to lysosomes

To explore the role of Ca^2+^ in maintaining lysosomal integrity, we looked at Ca^2+^ mediated changes in proteins on the lysosome. From earlier work we knew that GPN triggers appearance of ESCRTs on lysosomes (5). Among Ca^2+^-responsive proteins, ALG-2 is known to interact with and connect the ESCRTs TSG101, VPS37B/C, ALIX and IST1 (Fig. 4A) (21, 23–30). Given the elevation in Ca^2+^ induced by GPN, we wondered if ALG-2 might be responsible for recruiting ESCRTs to GPN-stressed lysosomes.

**Figure 4.**
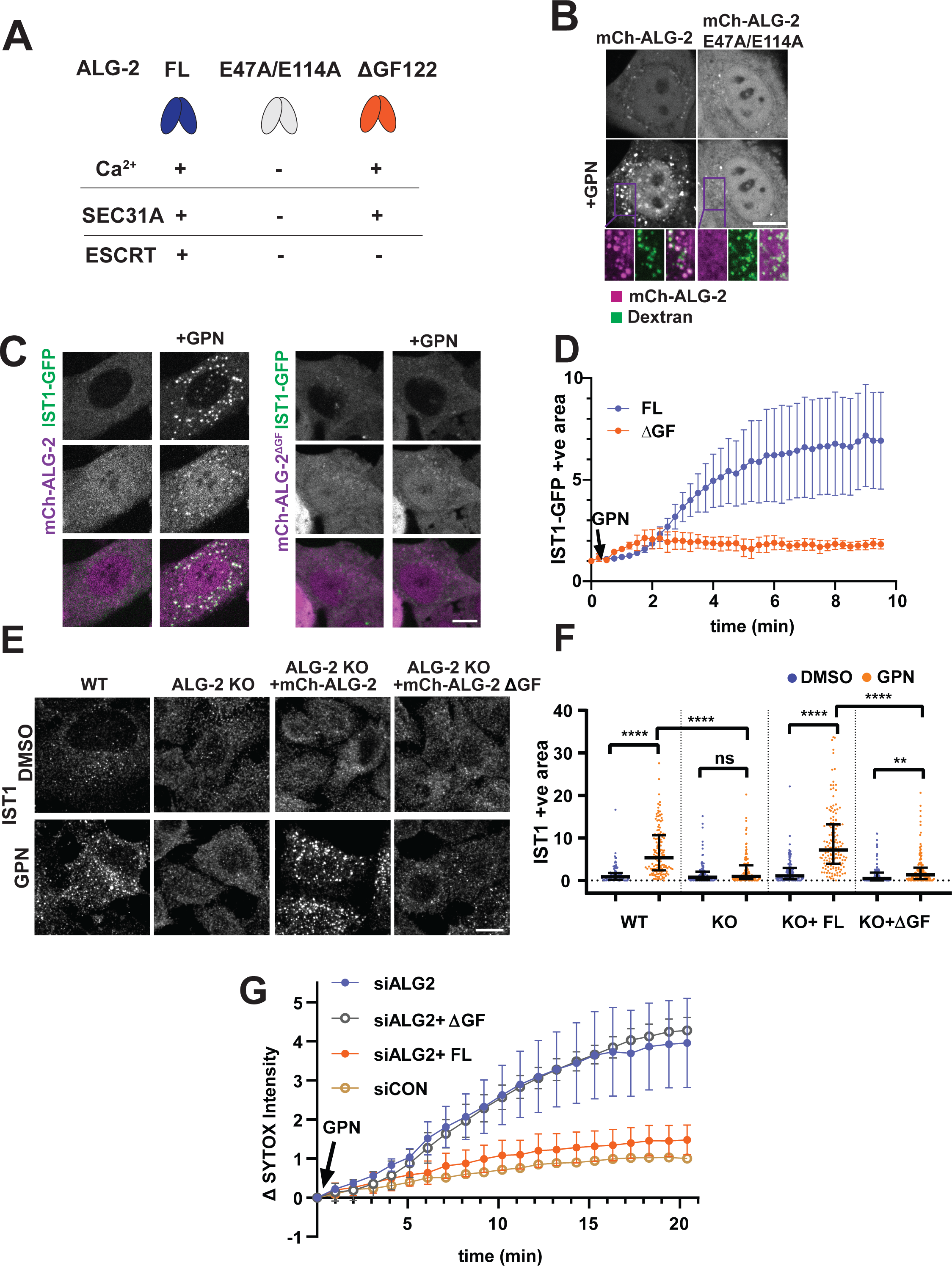
ALG-2 dependent ESCRT recruitment to lysosomes enhances resilience to osmotic stress. (A) Full length (FL) ALG-2 can bind Ca^2+^, SEC31A and ESCRTs. Mutants cannot as indicated by +/-. (B) ALG-2 KO HeLa cells were transfected with mCherry-ALG-2 or mCherry-ALG2^E47A/E114A^ and loaded with Oregon Green 70 kD dextran to label lysosomes. Cells were imaged at 1 min intervals before and after adding GPN (see also movie 2). (C) ALG-2 KO HeLa cells were co-transfected with IST1-GFP and mcherry-ALG-2(FL) or mCherry-ALG-2(ΔGF). Cells were imaged at 15 s intervals before and after adding GPN (see also Movie 3). (D) IST1-GFP positive area was normalized to the mean starting value for each group. Normalized area means ± SEM are plotted. More than 90 cells from 3 independent experiments for each condition were analyzed. (E) WT HeLa cells were transfected with mCherry, and ALG-2 KO HeLa cells were transfected with mCherry, mCherry-ALG-2 (FL) or mCherry-ALG-2^ΔGF122^ (ΔGF). Cells were treated with DMSO or GPN for 10 min and stained for IST1. IST1 signal above threshold was analyzed for each condition. (F) IST1 positive area in each group was normalized to the median value of IST1 positive area in WT DMSO group. Normalized median with interquartile range (n= 96, 142, 110, 116, 115, 145, 112 and 149 cells from 2 independent experiments) of positive area per cell are plotted. **** denotes P<0.0001; ** denotes P<0.01; ns denotes P>0.9999 by Kruskal– Wallis test. (G) HeLa cells were transfected with siCON or siALG-2. Cells transfected with siALG-2 were further transfected with siRNA resistant mCherry-ALG-2 (FL) or mCherry-ALG-2^ΔGF122^ (ΔGF). Cells were loaded with SYTOX and imaged live at 1 min intervals before and after adding GPN. Changes of nuclear SYTOX intensities (mean ± SEM, normalized to the mean value of nuclear SYTOX intensity change in siCON group at 20 min in each experiment; n>200 cells for each condition from 3 independent experiments) were measured and plotted for each time point. Kruskal–Wallis test was performed for the final time point. siALG-2 and siALG-2+ ΔGF were not significantly different (P>0.9999); siALG-2+FL and siCON were not significantly different (P>0.9999); siALG-2 and siCON were significantly different (P<0.05). Scale bars, 10 µm.

We began by looking for changes in ALG-2 localization following GPN treatment using live-cell spinning disc confocal microcopy. To avoid complications from coassembly with endogenous ALG-2, we transfected mCherry-ALG-2 into ALG-2 knockout (KO) HeLa cells (34). After adding GPN, mCherry-ALG-2 formed puncta that significantly overlapped with lysosomes marked by internalized 70kDa fluorescent dextran (Fig. 4B). Dimmer and transient puncta that did not overlap with dextran were also present (Fig. 4B, movie 2). These likely correspond to ER exit sites that engage ALG-2 (45, 46). A calcium binding mutant (E47A/E114A) of ALG-2 (Fig. 4A) (28, 47) did not redistribute in response to GPN (Fig. 4B).

To explore connections between ALG-2 and ESCRTs, we tracked the ESCRT-III protein IST1-GFP together with mCherry-ALG-2 in ALG-2 KO HeLa cells. After adding 0.2 mM GPN, IST1-GFP accumulated with mCherry-ALG-2 in puncta (Fig. 4C&D, movie 3). This did not happen with mCherry-ALG-2^ΔGF122^, a splice variant that lacks two amino acids and the ability to bind ESCRTs (Fig. 4A) (24, 25, 27), highlighting ALG-2’s role in bringing ESCRTs to GPN-stressed lysosomes.

To determine if ALG-2 is required to recruit ESCRTs in response to GPN, we compared changes in endogenous ESCRT proteins in WT and ALG-2 KO HeLa cells. GPN-induced IST1 accumulation in puncta was reduced in ALG-2 KO cells and readily restored by expressing mCherry-ALG-2 but not mCherry-ALG-2^ΔGF122^ (Fig. 4E&F). Similarly, GPN caused less accumulation of ALIX in puncta in ALG-2 KO cells (Fig. S8). Transiently depleting ALG-2 using siRNA also reduced ESCRT recruitment in response to GPN (Fig. S9). Together, these results establish that Ca^2+^-ALG-2 is essential for recruiting the ESCRT machinery following GPN exposure.

### ALG-2 enhances lysosomal resilience to osmotic stress

ALG-2 and interacting ESCRTs respond widely within cells treated with 0.2 mM GPN (Fig. 4), mirroring the pattern of juxta-lysosomal Ca^2+^ signals (Fig. 3). Given the contrast with the limited number of ruptured lysosomes (Fig. 2), we wondered if ALG-2 and the ESCRTs it recruits might play a protective role in the setting of osmotic stress. To test this idea, we examined the impact of ALG-2 depletion on lysosomal vulnerability to GPN as assessed by SYTOX leakage (Fig. 4G). In cells depleted of ALG-2 by siRNA (Fig. S9C), there was a striking ∼4-fold increase in nuclear SYTOX following addition of GPN. Expressing siRNA-resistant wild-type ALG-2 in these depleted cells maintained lysosomal integrity at levels comparable to control while expressing the ALG-2^ΔGF122^ splice variant did not (Fig. 4G). Thus by decreasing lysosomal rupture, the ALG-2-ESCRT axis increases lysosomal membrane resilience to osmotic stress.

To further assess the contribution of ALG-2 and associated ESCRTs to lysosomal membrane resilience, we extended our investigation to include the effects of chlorpromazine (CPZ). Using an intermediate concentration of 0.25 mM CPZ, based on its dose-dependent induction of GAL3 accumulation (Fig. S7A), we observed that ALG-2 depletion similarly resulted in increased lysosomal leakage (Fig. S7C). Thus, for both GPN and CPZ-induced stress, ALG-2 is a key player in maintaining lysosomal integrity.

### TRPML1 mediated Ca^2+^ release recruits ALG-2-ESCRT to lysosomes

Having established that GPN differentially elicits Ca^2+^ release and lysosomal rupture, we wondered whether juxta-lysosomal Ca^2+^ might itself suffice to recruit ALG-2 and ESCRTs or if there are additional damage-specific signals. Since ALG-2 responds to Ca^2+^ released by the lysosomal cation channel TRPML1 (34), we first tested the effect of TRPML1 activation on ESCRT recruitment. Using the TRPML1 agonist ML-SA5 we observed accumulation of ALIX-GFP in cytosolic puncta (Fig. 5A). Additional inclusion of thapsigargin (TG), a potent inhibitor of the ER Ca^2+^ ATPase that elevates juxta-ER Ca^2+^, led to more and brighter ALIX-GFP puncta (Fig. 5A) establishing that juxta-lysosomal Ca^2+^ indeed suffices to recruit the ESCRT machinery.

**Figure 5.**
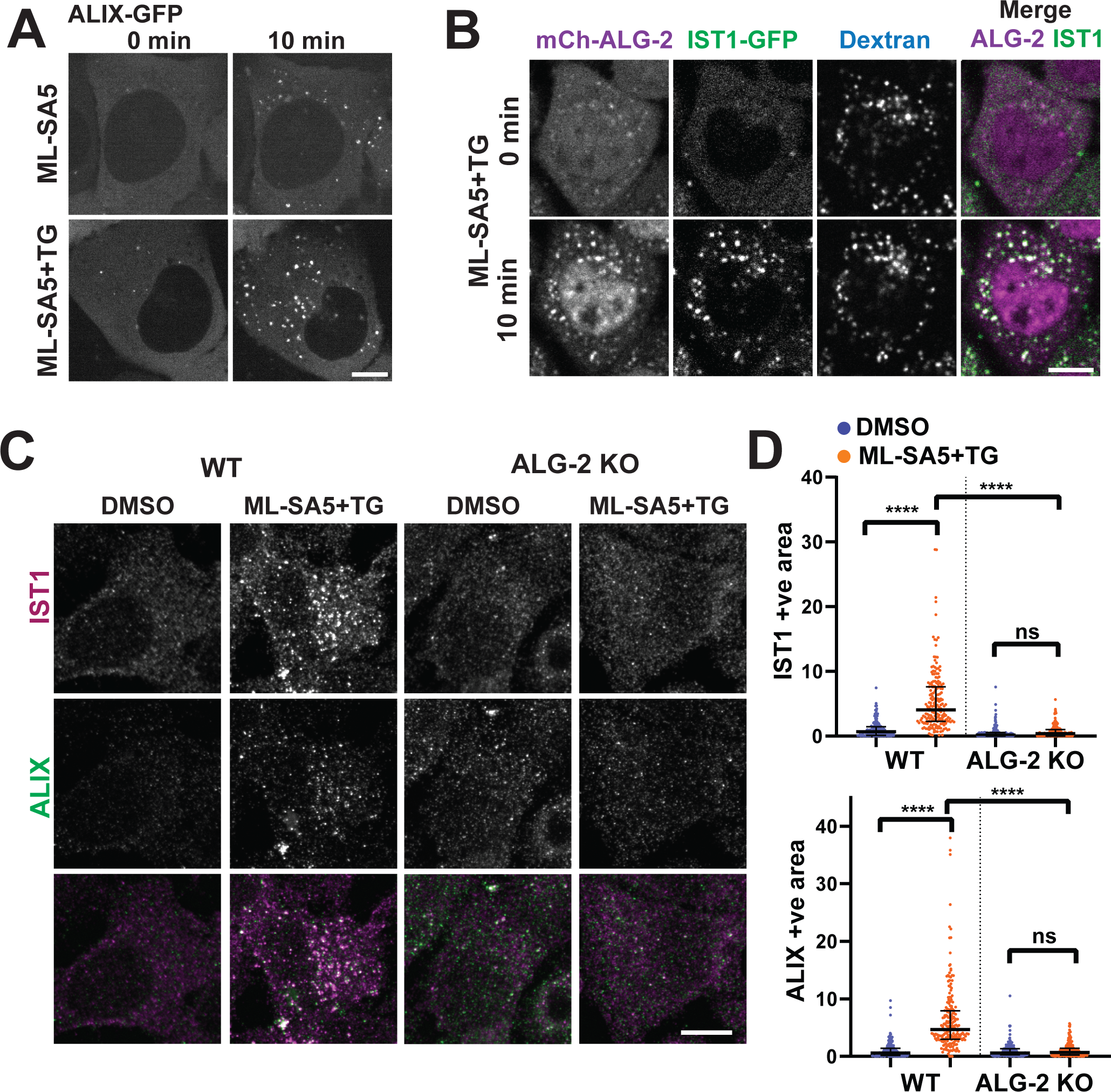
Lysosomal Ca^2+^ release triggers ALG-2 dependent ESCRT recruitment without membrane damage. (A) U2OS cells transfected with ALIX-GFP were treated with ML-SA5 or ML-SA5+TG for 10 min and imaged live before and after treatment. (B) ALG-2 KO HeLa cells co-transfected with mCherry-ALG-2 and IST1-GFP were loaded with AF647 dextran to label lysosomes. Cells were imaged live before and after ML-SA5 with TG co-treatment. (C) WT and ALG-2 KO HeLa cells were treated with DMSO or ML-SA5 with TG for 10 min and co-stained for IST1 and ALIX. IST1 and ALIX positive area were analyzed in each condition. (D) The ESCRT positive area in each group was normalized to the median value of positive area in WT DMSO group for both IST1 and ALIX. Normalized median with interquartile range (more than 150 cells from 2 independent experiments for each condition) of IST1 and ALIX positive area per cell are plotted. **** denotes P<0.0001; ns denotes P= 0.0668 in IST1 and P>0.9999 in ALIX by Kruskal–Wallis test. Scale bars, 10µm.

To examine the role of ALG-2 in this Ca^2+^ triggered ESCRT recruitment, we looked at ALG-2 and ESCRT dynamics in ALG-2 KO HeLa cells. In ALG-2 KO cells overexpressing mCherry-ALG-2 and IST1-GFP, combined addition of ML-SA5 and TG triggered assembly of bright IST1-GFP and mCherry-ALG-2 containing puncta (Fig. 5B). ALG-2^ΔGF122^ did not support IST1-GFP recruitment (Fig. S10). To exclude off-target effects of ML-SA5, we also tested a second, structurally distinct, TRPML1 agonist, ML-SA1(14). Like ML-SA5, ML-SA1 combined with TG promoted co-assembly of mCherry-ALG-2 and IST1-GFP on punctate lysosomes (Fig. S10). To exclude off-target effects of TG, we used the Ca^2+^ ionophore Ionomycin to raise intracellular Ca^2+^ and found that it also triggered IST1-GFP puncta formation in conjunction with ML-SA5 but not alone (Fig. S10).

We next surveyed the effects of Ca^2+^ mobilization on endogenous ESCRT proteins in WT and ALG-2 KO HeLa cells. Immunostaining IST1, ALIX and CHMP3 showed redistribution upon treatment with ML-SA5 and TG in WT but not ALG-2 KO cells (Fig 5C, D and Fig. S11). We confirmed that these Ca^2+^ mobilizing agents did not themselves cause lysosomal damage, as there was no change in nuclear SYTOX intensity (Fig. S12). Altogether, we conclude that local Ca^2+^ signals bring ESCRTs together on lysosomes in an ALG-2 dependent manner that does not require a membrane damage-specific signal.

### TRPML1 activation protects lysosomes from osmotic stress

Finally, since GPN-triggered Ca^2+^ release increases lysosomal membrane resilience, we wondered whether channel-mediated lysosomal Ca^2+^ release would similarly precondition lysosomes to make them less sensitive to osmotic stress. Brief pretreatment with ML-SA5 alone or together with TG significantly reduced SYTOX release upon addition of 0.2 mM GPN while pretreatment with TG alone failed to offer similar protection (Fig. 6A). This is consistent with a Ca^2+^-driven protective ESCRT response on lysosomes.

**Figure 6.**
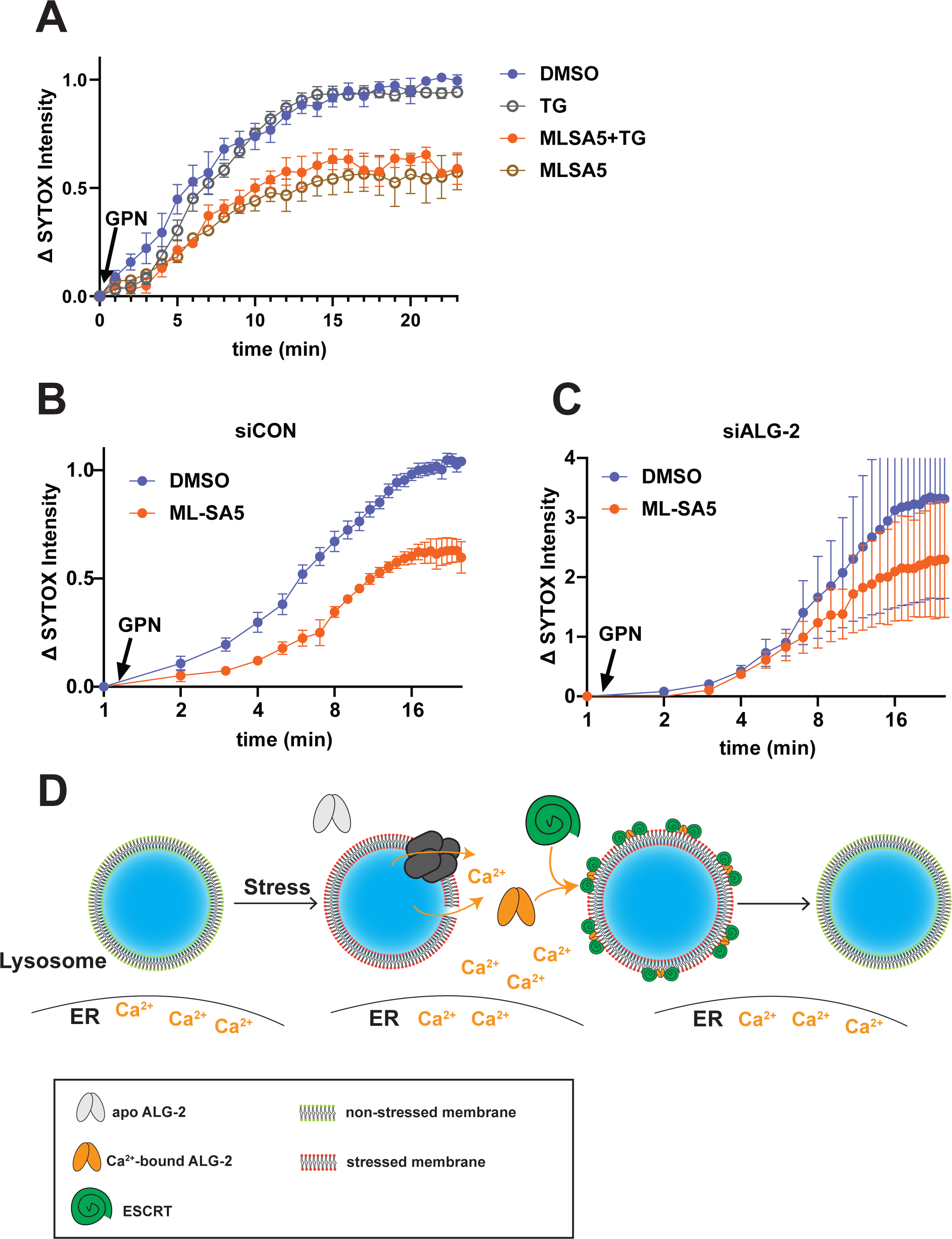
TRPML1 activation confers resilience to lysosomal osmotic stress. (A) HeLa cells loaded with SYTOX were incubated with indicated treatments for 3min prior to addition of GPN. Cells were then imaged at 1 min intervals during exposure to GPN. Nuclear SYTOX intensity changes (mean ± SEM, normalized the mean value of CTRL group at 25 min; n>200 cells for each condition from at least 3 independent experiments) were measured and plotted for each time point. Kruskal–Wallis test was performed for the final time point. DMSO and TG were not significantly different (P>0.9999); ML-SA5+TG and MLSA-5 were not significantly different (P>0.9999); DMSO and ML-SA5 were significantly different (P<0.05). HeLa cells transfected with siCON (B) and siALG-2 (C) were loaded with SYTOX and pre-incubated with DMSO or ML-SA5 for 3min. Cells were imaged live at 1 min intervals during exposure to GPN. Nuclear SYTOX intensity changes (mean ± SEM, normalized to the mean value of siCON CTRL group at 20 min; n>200 cells for each condition from at least 3 independent experiments) were measured and plotted in (B&C) for each time point. Unpaired T-tests were performed for each time point in both siCON and siALG-2 groups. In siCON group, DMSO and ML-SA5 were significantly different starting from 1 min after GPN (1-4 min: P<0.05; 5-10 min: P<0.01; 6-17 min: P<0.001; after 18 min: P<0.0001). In siALG-2 group, DMSO and ML-SA5 were not significantly different from 0-17 min (P>0.05). (D) Schematic of Ca^2+^-ALG-2-ESCRT mediated membrane resilience. Stress-induced Ca^2+^ released via lysosomal leakage or activation of a Ca^2+^-permeable channel preconditions ALG-2-ESCRT assembly on lysosomes to protect the organelle from rupture.

To directly assess contributions of ALG-2 and associated ESCRTs to lysosomal membrane resilience, we compared GPN-induced SYTOX release in wild-type and ALG-2 depleted cells following TRPML1 activation. The protective effect of ML-SA5 during the first several minutes of exposure to GPN was largely abolished by loss of ALG-2 (Fig. 6B&C). At later stages, ML-SA5 pretreatment had some albeit weaker protective effect in ALG-2 depleted cells. Altogether, these observations confirm that lysosomal Ca^2+^ release enhances resilience to osmotic membrane damage at least in part by engaging ALG-2 and ESCRTs.

## Discussion

Our findings establish that Ca^2+^, ALG-2 and ESCRTs contribute to lysosomal membrane resilience in the face of acute osmotic stress (Fig. 6D). Ca^2+^ released from osmotically challenged lysosomes and the adjacent ER triggers rapid ALG-2 dependent ESCRT accumulation, enhancing the ability of lysosomal membranes to withstand osmotic pressure and thus avoid organelle rupture. Elevating juxta-lysosomal Ca^2+^ by activating the cation-permeable TRPML1 channel similarly recruits ALG-2 and ESCRTs to protect lysosomes. This preventative role for ALG-2 and associated ESCRTs adds new insight into how the broadly deployed ESCRT machinery contributes to lysosomal membrane homeostasis.

Despite GPN being commonly used to release and deplete lysosomal Ca^2+^ stores (13–18), it has not been widely deployed in cellular studies of lysosomal membrane permeabilization. Responses to GPN have instead served – without the benefit of much scrutiny – to confirm engagement of factors identified using LLOMe (L-Leucyl-L-Leucine O-methyl ester) (5–7). LLOMe permeabilizes lysosomes via a detergent-like mechanism in which small-molecule leakage precedes full membrane rupture (48, 49), which as shown in Fig. 2 differs from the simultaneous loss of small and large molecules caused by GPN. Interestingly, ALG-2 accumulates on lysosomes damaged with LLOMe (5, 6, 23) but, in contrast to what we find with GPN, is not required to recruit ESCRTs (35). This divergence highlights differences in the membrane damage and cellular responses elicited by these reagents and the need for more nuanced approaches to dissecting cellular responses to lysosomal perturbation.

Our observation that moderate concentrations of GPN trigger lysosomal Ca^2+^ release without significant SYTOX leakage may explain differing conclusions about the effects of GPN in previous studies (19, 20, 40). It also raises questions about how Ca^2+^ escapes the lysosome in the setting of non-rupturing osmotic stress. Ca^2+^ could traverse transient and size-limited defects in the membrane or escape via stress-sensitive Ca^2+^-permeant channels such as TRPMLs or TPCs. Interestingly, TRPML1 is responsive to oxidative stress and starvation (41, 50) and could be acting to limit lysosomal membrane damage. Additionally, TRPML2 is activated by osmotic pressure and high endomembrane tension (51) and the lipid kinase PIKfyve is stimulated by osmotic stress to generate channel-activating PI(3,5)P_2_ (52–54). Further work will be needed to establish whether these or other mechanisms underlie protective lysosomal Ca^2+^ signaling.

The role of Ca^2+^-ALG-2 and the ESCRTs it engages extends beyond the lysosome to the plasma membrane where ALG-2 was first reported to accumulate at sites of damage and contribute to their repair (21, 55). While prevailing models for ESCRT-dependent membrane repair focus on ESCRT-mediated vesicle formation and release, the fact that Ca^2+^-ALG-2 also responds and recruits ESCRTs to plasma membranes treated with Ca^2+^ ionophore (21) raises the possibility of a broader, damage-independent, role in enhancing membrane resilience. The overall concept that ESCRTs could be important in maintaining membrane integrity is further supported by recent studies showing that ESCRT-III proteins extend into all kingdoms of life. A key component of the bacterial phage-shock-protein response pathway, PspA, known for its role in stabilizing the bacterial inner membrane, shares a common structure with the ESCRT-III superfamily (56, 57). Similarly, the chloroplast protein VIPP1, also an ESCRT-III protein (56), maintains chloroplast envelope and thylakoid membrane integrity under environmental stress (58–62).

In summary, previous studies have established that the ESCRT machinery repairs lysosomes as they recover from membrane damage. Here, we demonstrate that ESCRTs also have a proactive role in fortifying lysosomal membranes against stress. This suggests that ESCRTs operate in a dynamic and potentially multi-faceted fashion to ensure lysosomal integrity, highlighting the machinery’s essential function in cellular resilience to environmental and physiologic stress.

## Materials and methods

### Reagents

Compounds were used at the following final concentrations unless explicitly stated: 1 μM SYTOX Green (S7020, Invitrogen); 200 μM GPN (no. sc-252858; Santa Cruz Biotechnology); 25 μM BAPTA-AM (no. 15551; Cayman Chemical); 200 μM E64d (no. 13533; Cayman Chemical); 2 μM ML-SA5 (No. AOB11116, Aobious); 25 μM ML-SA1 (No. 4746, Tocris Bioscience); 2.5 μM Thapsigargin (AC328570010, Thermo Scientific); 5 μM Ionomycin (No. 10004974, Cayman Chemical); 250 μM CPZ (No. 16129, Cayman Chemical); 0.5 mg/mL Alexa Fluor 647 10,000 MW Dextran (D22914, Invitrogen); 0.15mg/mL Oregon Green 70,000 MW Dextran (D7173, Invitrogen).

The mammalian expression vector encoding mCherry-GAL3 has been described(5). pCMV R-CEPIA1er was a gift from M. Iino (Addgene plasmid # 58216). The plasmid encoding TMEM192-GCAMP7 was constructed by cloning TMEM192 and GCAMP7 sequence into pcDNA 3.1/Zeo (+) using Gibson Assembly (E2611, NEB). The plasmids encoding GCAMP7-TRPML1, mCherry-ALG-2^ΔGF122^ and mCherry-ALG2^E47A/E114A^ were generously supplied by H. Xu at University of Michigan and have been described(34). siRNA resistant mCherry-ALG-2 was created by mutagenesis (primers sequences 5’-ATTTCGATGTTTGACCGTGAGAAC-3’, 5’-AATGGACCTGACAGTCACTGGATTAAA-3’) using a Q5 Site-Directed Mutagenesis Kit (E0554S, NEB). The plasmid encoding IST1-GFP is as described (63) and that encoding ALIX-H30-GFP as described (64) was a kind gift from S. Saffarian (Univ. of Utah).

### Cell culture

All cells were maintained at 37°C and supplemented with 5% CO2. HeLa human cervical adenocarcinoma, U2OS human osteosarcoma originally from the American Type Culture Collection (ATCC; Manassas, VA, USA) were grown in Dulbecco’s modified Eagle’s medium (DMEM) (no. 11965-084; Gibco, Carlsbad, CA, USA) supplemented with 9% v/v fetal bovine serum (FBS; Atlanta Biologicals, S11150). ALG-2 KO HeLa cells were a gift from H. Xu at University of Michigan and have been described(34).

### RNA interference

Subconfluent cells were suspended by trypsinization and transfected using Lipofectamine RNAiMAX Transfection Reagent (Invitrogen, no. 13778150) according to the manufacturer’s instructions. The final concentration was 15 nM total siRNA and 0.3% v/v transfection reagent. The cell suspension was dispensed into a well of a six-well plate and incubated as described above. After 48 to 72 hours, cells were suspended by trypsinization, seeded at an appropriate density, and incubated for an additional 16 to 24 hours. The siRNA sequences used were against human ALG-2 (5’-GGUCGAUCAUAUCCAUGUUdTdT-3’).

### DNA transfection

Cells were suspended by trypsinization and transfected using Lipofectamine 2000 (Invitrogen, no. 11668027) according to the manufacturer’s instructions. The final concentration was 1.5 μg/ml total DNA, and 0.2% v/v transfection reagent. The cell suspension was dispensed into a culture vessel appropriate for the intended application. Medium was replaced after 6h and transfected cells were further incubated for 16 to 24 hours before use.

### Confocal microscopy

Images were acquired on an Olympus IX83 microscope with a Yokogawa CSU-W1 spinning disk confocal scanner unit, an Olympus PlanApo 60x 1.42 NA objective, and an ImageEMX2 EMCCD digital camera or Hamamatsu Orca Fusion CMOS digital camera. Colors were imaged as follows: Blue, 405 nm laser, 460/50 emission filter; Green, 488 nm laser, 525/50 emission filter, Red, 561 nm laser, 605/52 emission filter and Far Red, 638 nm laser, 700/75 emission filter. Images were acquired with Metamorph software Version 7.10.3279 (Molecular Devices).

### Live imaging

Cells were seeded in a four-chamber, no. 1.5 glass-bottom dish (Cellvis, D35C4-20-1.5-N) and cultured as appropriate for the intended experiment. Before imaging, medium was replaced with warm imaging solution (DMEM, high glucose, HEPES, no phenol red (Gibco, 21063029) supplemented with 10% v/v fetal bovine serum). The dish was immediately transferred to a Tokai Hit (model: STXG-WSKMX-SET) stage-top incubator preheated to 37°C. The dish was allowed to equilibrate for 10 min before initiating acquisition. Imaging solution intended for drug treatments was warmed to 37°C.

### SYTOX leak assay

SYTOX loading: Subconfluent cells were suspended by trypsinization and diluted in growth medium, and the cell suspension was mixed with 1 μM SYTOX and dispensed into four-well glass-bottom dish. Alternately, cells were seeded at appropriate density in four-well glass-bottom dish and allowed to adhere for 6-12 hours, after which medium was then replaced with fresh growth medium containing 1 μM SYTOX.

SYTOX chase: After 16 hours, medium was replaced with fresh medium and cells were incubated for a further 4-6 hours to allow the SYTOX to accumulate in late endosomes and lysosomes.

SYTOX imaging: Before imaging, 300 μL imaging solution was added to each well. Desired reagents were diluted in 300 μL warm imaging solution at 2x of the final concentration for each well. During live imaging, desired reagent treatment was applied at indicated time point.

### Ca^2+^ imaging

U2OS cells transfected with plasmid encoding CEPIAer, TMEM192-GCAMP7 and/or GCAMP7-TRPML1 were seeded in four-well glass-bottom dish. Before imaging, medium was replaced with warm imaging solution or zero Ca^2+^ solution (145 mM NaCl, 5 mM KCl, 3 mM MgCl_2_, 10 mM glucose, 1 mM EGTA, and 20 mM HEPES). During live imaging, desired reagent treatment was applied at indicated time point.

### Immunofluorescence

Cells grown over no. 1.5 round glass coverslips (Electron Microscopy Sciences) were fixed in 4% w/v paraformaldehyde (Electron Microscopy Sciences) in phosphate-buffered saline (PBS) for 15 min at room temperature, rinsed with PBS, and permeabilized in 0.1% v/v Triton X-100 (no. 28314; Pierce Biotechnology). Cells were immunolabeled in blocking solution for 1 hour at RT with antibodies against the following proteins: ALIX at 5 μg/ml (mouse; no. 634502, Biolegend); IST1 at 0.4 μg/ml (rabbit; no. 19842-1-AP, ProteinTech); CHMP3 at 1 μg/ml (mouse; no. sc-166361, Santa Cruz). After rinsing with PBS, goat secondary antibodies conjugated to Alexa Fluor 488, 555, or 647 fluorescent dyes (Molecular Probes, Carlsbad, CA, USA) were diluted to 1 μg/ml in blocking solution were added cells for 45 min. After rinsing with PBS, 0.25 μg/ml of 4′,6-diamidino-2-phenylindole (DAPI; Molecular Probes) was added to immunolabeled cells for 10 min. After rinsing with PBS, coverslips were mounted with homemade Gelvatol and allowed to cure for 24h.

### Image analysis

All image analyses were performed in ImageJ. In general, the intensities or protein positive areas were normalized to the control group in each experiment. Details can be found in corresponding figure legends. For SYTOX leak analyses, nuclear SYTOX intensity was measured for each frame by manually placing a circular ROI in the center of each nucleus. The change in intensity was calculated by subtracting the first frame value from each subsequent value. For Ca^2+^ imaging, individual cells were circled and Ca^2+^ indicator intensities were measured. ΔF/F_0_ was calculated and plotted for each time point. Off-cell background was subtracted from F_0_. For ESCRT positive area analyses, a threshold was empirically determined, and the same threshold was applied to all images analyzed within each experiment. The ImageJ command Analyze Particle was used to measure the above threshold protein positive area per cell.

### Statistics

Statistical significance was calculated using GraphPad Prism 8, performing Kruskal-Wallis one-way unpaired ANOVA statistical tests or nonparametric unpaired t-tests that did not assume a Gaussian distribution.

## Acknowledgments

We thank Kim Martin for help with Galectin-3 GPN dosage experiments, Kimberly Edicha for help with molecular cloning, and members of the Hanson lab for discussion and feedback. This research was supported by NIH R01 GM122434 (to P.I.H.).

---

**Supplemental Movie 1.** Whole field timelapse recording that corresponds to the time series shown in Fig. 1B. GPN was added to U2OS cell with SYTOX-loaded lysosomes at indicated time point. Images were captured every 15s.

**Supplemental Movie 2.** Timelapse recording that corresponds to the time series shown in Fig. 4B. GPN was added at indicated time to ALG-2 KO HeLa cells transfected with mCherry-ALG-2 or mCherry-ALG2^E47A/E114A^ and loaded with Oregon Green 70 kD dextran to label lysosomes. Images were captured every 1 min.

**Supplemental Movie 3.** Timelapse recording that corresponds to the time series shown in Fig. 4C. GPN was added at indicated time to ALG-2 KO HeLa cells co-transfected with IST1-GFP and mcherry-ALG-2(FL) or ΔGF122. Images were captured every 15 s.

## References

1. I. Paz et al., Galectin-3, a marker for vacuole lysis by invasive pathogens. Cell Microbiol 12, 530–544 (2010).

2. M. Jadot, C. Colmant, S. Wattiaux-De Coninck, R. Wattiaux, Intralysosomal hydrolysis of glycyl-L-phenylalanine 2-naphthylamide. Biochem J. 219, 965–70 (1984).

3. G. N. Joshi, A. M. Goetjen, D. A. Knecht, Silica particles cause NADPH oxidase–independent ROS generation and transient phagolysosomal leakage. Molecular Biology of the Cell 26, 3150–3164 (2015).

4. V. Hornung et al., Silica crystals and aluminum salts activate the NALP3 inflammasome through phagosomal destabilization. Nature Immunology 9, 847–856 (2008).

5. M. L. Skowyra, P. H. Schlesinger, T. V. Naismith, P. I. Hanson, Triggered recruitment of ESCRT machinery promotes endolysosomal repair. Science 360 (2018).

6. M. Radulovic et al., ESCRT-mediated lysosome repair precedes lysophagy and promotes cell survival. EMBO J 37 (2018).

7. J. X. Tan, T. Finkel, A phosphoinositide signalling pathway mediates rapid lysosomal repair. Nature 609, 815–821 (2022).

8. P. Niekamp et al., Ca2+-activated sphingomyelin scrambling and turnover mediate ESCRT-independent lysosomal repair. Nature Communications 13, 1875 (2022).

9. R. Goldman, A. Kaplan, Rupture of rat liver lysosomes mediated by l-amino acid esters. Biochimica et Biophysica Acta (BBA) - Biomembranes 318, 205–216 (1973).

10. J. P. Reeves, Accumulation of amino acids by lysosomes incubated with amino acid methyl esters. Journal of Biological Chemistry 254, 8914–8921 (1979).

11. D. L. Thiele, P. E. Lipsky, The action of leucyl-leucine methyl ester on cytotoxic lymphocytes requires uptake by a novel dipeptide-specific facilitated transport system and dipeptidyl peptidase I-mediated conversion to membranolytic products. Journal of Experimental Medicine 172, 183–194 (1990).

12. T. O. Berg, E. Strømhaug, T. Løvdal, O. Seglen, T. Berg, Use of glycyl-L-phenylalanine 2-naphthylamide, a lysosome-disrupting cathepsin C substrate, to distinguish between lysosomes and prelysosomal endocytic vacuoles. Biochem J. 300, 229–36 (1994).

13. T. Haller, P. Dietl, P. Deetjen, H. Völkl, The lysosomal compartment as intracellular calcium store in MDCK cells: a possible involvement in InsP3-mediated Ca2+ release. Cell Calcium 19, 157–165 (1996).

14. D. Shen et al., Lipid storage disorders block lysosomal trafficking by inhibiting a TRP channel and lysosomal calcium release. Nature Communications 3, 731 (2012).

15. A. G. Garrity et al., The endoplasmic reticulum, not the pH gradient, drives calcium refilling of lysosomes. Elife 5 (2016).

16. B. S. Kilpatrick, E. R. Eden, A. H. Schapira, C. E. Futter, S. Patel, Direct mobilisation of lysosomal Ca2+ triggers complex Ca2+ signals. Journal of Cell Science 126, 60–66 (2013).

17. G. C. Churchill et al., NAADP Mobilizes Ca2+ from Reserve Granules, Lysosome-Related Organelles, in Sea Urchin Eggs. Cell 111, 703–708 (2002).

18. X. Zhang et al., Rapamycin directly activates lysosomal mucolipin TRP channels independent of mTOR. PLOS Biology 17, e3000252 (2019).

19. A. J. Morgan, Y. Yuan, S. Patel, A. Galione, Does lysosomal rupture evoke Ca2+ release? A question of pores and stores. Cell Calcium 86, 102139 (2020).

20. Y. Yuan et al., The lysosomotrope GPN mobilises Ca2+ from acidic organelles. Journal of Cell Science 134, jcs256578 (2021).

21. L. L. Scheffer et al., Mechanism of Ca(2)(+)-triggered ESCRT assembly and regulation of cell membrane repair. Nat Commun 5, 5646 (2014).

22. M. Maki, Structures and functions of penta-EF-hand calcium-binding proteins and their interacting partners: enigmatic relationships between ALG-2 and calpain-7. Biosci Biotechnol Biochem 84, 651–660 (2020).

23. S. Shukla, K. P. Larsen, C. Ou, K. Rose, J. H. Hurley, In vitro reconstitution of calcium-dependent recruitment of the human ESCRT machinery in lysosomal membrane repair. Proceedings of the National Academy of Sciences 119, e2205590119 (2022).

24. M. Missotten, K. Nichols A Fau - Rieger, R. Rieger K Fau - Sadoul, R. Sadoul, Alix, a novel mouse protein undergoing calcium-dependent interaction with the apoptosis-linked-gene 2 (ALG-2) protein. Cell Death Differ. 6,124-9. (1999).

25. K. Katoh et al., The penta-EF-hand protein ALG-2 interacts directly with the ESCRT-I component TSG101, and Ca2+-dependently co-localizes to aberrant endosomes with dominant-negative AAA ATPase SKD1/Vps4B. Biochem J 391, 677–685 (2005).

26. M. Okumura, T. Takahashi, H. Shibata, M. Maki, Mammalian ESCRT-III-related protein IST1 has a distinctive met-pro repeat sequence that is essential for interaction with ALG-2 in the presence of Ca2+. Biosci Biotechnol Biochem 77, 1049–1054 (2013).

27. S. Tarabykina, A. L. Moller, I. Durussel, J. Cox, M. W. Berchtold, Two forms of the apoptosis-linked protein ALG-2 with different Ca(2+) affinities and target recognition. J Biol Chem 275, 10514–10518 (2000).

28. H. Suzuki et al., Structural Basis for Ca2+-Dependent Formation of ALG-2/Alix Peptide Complex: Ca2+/EF3-Driven Arginine Switch Mechanism. Structure 16, 1562–1573 (2008).

29. M. Okumura et al., Penta-EF-hand protein ALG-2 functions as a Ca2+-dependent adaptor that bridges Alix and TSG101. Biochem Biophys Res Commun 386, 237–241 (2009).

30. M. Okumura, A. M. Katsuyama, H. Shibata, M. Maki, VPS37 isoforms differentially modulate the ternary complex formation of ALIX, ALG-2, and ESCRT-I. Biosci Biotechnol Biochem 77, 1715-1721 (2013).

31. J. M. la Cour et al., ALG-2 participates in recovery of cells after plasma membrane damage by electroporation and digitonin treatment. PLOS ONE 13, e0204520 (2018).

32. J. Westman et al., Calcium-dependent ESCRT recruitment and lysosome exocytosis maintain epithelial integrity during Candida albicans invasion. Cell Rep 38, 110187 (2022).

33. S. Vergarajauregui, J. A. Martina, R. Puertollano, Identification of the penta-EF-hand protein ALG-2 as a Ca2+-dependent interactor of mucolipin-1. J Biol Chem 284, 36357–36366 (2009).

34. X. Li et al., A molecular mechanism to regulate lysosome motility for lysosome positioning and tubulation. Nat Cell Biol 18, 404–417 (2016).

35. W. W. Yim, H. Yamamoto, N. Mizushima, Annexins A1 and A2 are recruited to larger lysosomal injuries independently of ESCRTs to promote repair. FEBS Lett 596, 991–1003 (2022).

36. B. L. Roth, M. Poot, S. T. Yue, P. J. Millard, Bacterial viability and antibiotic susceptibility testing with SYTOX green nucleic acid stain. Appl Environ Microbiol 63, 2421–2431 (1997).

37. N. A. Bright, L. J. Davis, J. P. Luzio, Endolysosomes Are the Principal Intracellular Sites of Acid Hydrolase Activity. Current Biology 26, 2233–2245 (2016).

38. L.-K. Roni et al., SLC12A9 is a lysosome-detoxifying ammonium – chloride co-transporter. bioRxiv 10.1101/2023.05.22.541801, 2023.2005.2022.541801 (2023).

39. S. Aits et al., Sensitive detection of lysosomal membrane permeabilization by lysosomal galectin puncta assay. Autophagy 11, 1408–1424 (2015).

40. P. Atakpa, L. M. van Marrewijk, M. Apta-Smith, S. Chakraborty, C. W. Taylor, GPN does not release lysosomal Ca(2+) but evokes Ca(2+) release from the ER by increasing the cytosolic pH independently of cathepsin C. J Cell Sci 132 (2019).

41. X. Zhang et al., MCOLN1 is a ROS sensor in lysosomes that regulates autophagy. Nature Communications 7, 12109 (2016).

42. M. Ohkura et al., Genetically Encoded Green Fluorescent Ca2+ Indicators with Improved Detectability for Neuronal Ca2+ Signals. PLOS ONE 7, e51286 (2012).

43. J. Suzuki et al., Imaging intraorganellar Ca2+ at subcellular resolution using CEPIA. Nature Communications 5, 4153 (2014).

44. L. Yu et al., Small-molecule activation of lysosomal TRP channels ameliorates Duchenne muscular dystrophy in mouse models. Science Advances 6, eaaz2736 (2020).

45. H. Shibata, H. Suzuki, H. Yoshida, M. Maki, ALG-2 directly binds Sec31A and localizes at endoplasmic reticulum exit sites in a Ca2+-dependent manner. Biochem Biophys Res Commun 353, 756–763 (2007).

46. S. Shukla et al., Mechanism and cellular function of direct membrane binding by the ESCRT and ERES-associated Ca2+-sensor ALG-2. Proceedings of the National Academy of Sciences 121, e2318046121 (2024).

47. K. W. H. Lo, Q. Zhang, M. Li, M. Zhang, Apoptosis-Linked Gene Product ALG-2 Is a New Member of the Calpain Small Subunit Subfamily of Ca2+-Binding Proteins. Biochemistry 38, 7498–7508 (1999).

48. D. L. Thiele, P. E. Lipsky, Mechanism of L-leucyl-L-leucine methyl ester-mediated killing of cytotoxic lymphocytes: dependence on a lysosomal thiol protease, dipeptidyl peptidase I, that is enriched in these cells. Proceedings of the National Academy of Sciences 87, 83–87 (1990).

49. U. Repnik et al., L-leucyl-L-leucine methyl ester does not release cysteine cathepsins to the cytosol but inactivates them in transiently permeabilized lysosomes. J Cell Sci 130, 3124–3140 (2017).

50. W. Wang et al., Up-regulation of lysosomal TRPML1 channels is essential for lysosomal adaptation to nutrient starvation. Proceedings of the National Academy of Sciences 112, E1373–E1381 (2015).

51. C.-C. Chen et al., TRPML2 is an osmo/mechanosensitive cation channel in endolysosomal organelles. Science Advances 6, eabb5064 (2020).

52. N. Jin, Y. Jin, L. S. Weisman, Early protection to stress mediated by CDK-dependent PI3,5P2 signaling from the vacuole/lysosome. Journal of Cell Biology 216, 2075–2090 (2017).

53. X.-p. Dong et al., PI(3,5)P2 controls membrane trafficking by direct activation of mucolipin Ca2+ release channels in the endolysosome. Nature Communications 1, 38 (2010).

54. X. Wang et al., TPC Proteins Are Phosphoinositide-Activated Sodium-Selective Ion Channels in Endosomes and Lysosomes. Cell 151, 372–383 (2012).

55. S. L. Sønder et al., Annexin A7 is required for ESCRT III-mediated plasma membrane repair. Scientific Reports 9, 6726 (2019).

56. J. Liu et al., Bacterial Vipp1 and PspA are members of the ancient ESCRT-III membrane-remodeling superfamily. Cell 184, 3660–3673.e3618 (2021).

57. B. Junglas et al., PspA adopts an ESCRT-III-like fold and remodels bacterial membranes. Cell 184, 3674–3688.e3618 (2021).

58. L. Zhang, Y. Kato, S. Otters, U. C. Vothknecht, W. Sakamoto, Essential Role of VIPP1 in Chloroplast Envelope Maintenance in Arabidopsis. The Plant Cell 24, 3695–3707 (2012).

59. A. Nordhues et al., Evidence for a Role of VIPP1 in the Structural Organization of the Photosynthetic Apparatus in Chlamydomonas. The Plant Cell 24, 637–659 (2012).

60. L. Zhang, H. Kondo, H. Kamikubo, M. Kataoka, W. Sakamoto, VIPP1 Has a Disordered C-Terminal Tail Necessary for Protecting Photosynthetic Membranes against Stress Plant Physiology 171, 1983-1995 (2016).

61. N. Ohnishi, L. Zhang, W. Sakamoto, VIPP1 Involved in Chloroplast Membrane Integrity Has GTPase Activity in Vitro. Plant Physiology 177, 328–338 (2018).

62. T. K. Gupta et al., Structural basis for VIPP1 oligomerization and maintenance of thylakoid membrane integrity. Cell 184, 3643–3659.e3623 (2021).

63. K. C. Amy et al., IST1 regulates select recycling pathways. bioRxiv 10.1101/2023.07.31.551359, 2023.2007.2031.551359 (2023).

64. P.-I. Ku, M. Bendjennat, J. Ballew, M. B. Landesman, S. Saffarian, ALIX Is Recruited Temporarily into HIV-1 Budding Sites at the End of Gag Assembly. PLOS ONE 9, e96950 (2014).

