## Supplemental Material for "Ca^2+^-sensor ALG-2 engages ESCRTs to enhance lysosomal membrane resilience to osmotic stress"

### Supplementary Materials

**This file includes:**
**Figs. S1 to S12**
**Captions for Movies 1-3**

**Other Supplementary Materials for this manuscript include Movie 1-3.**

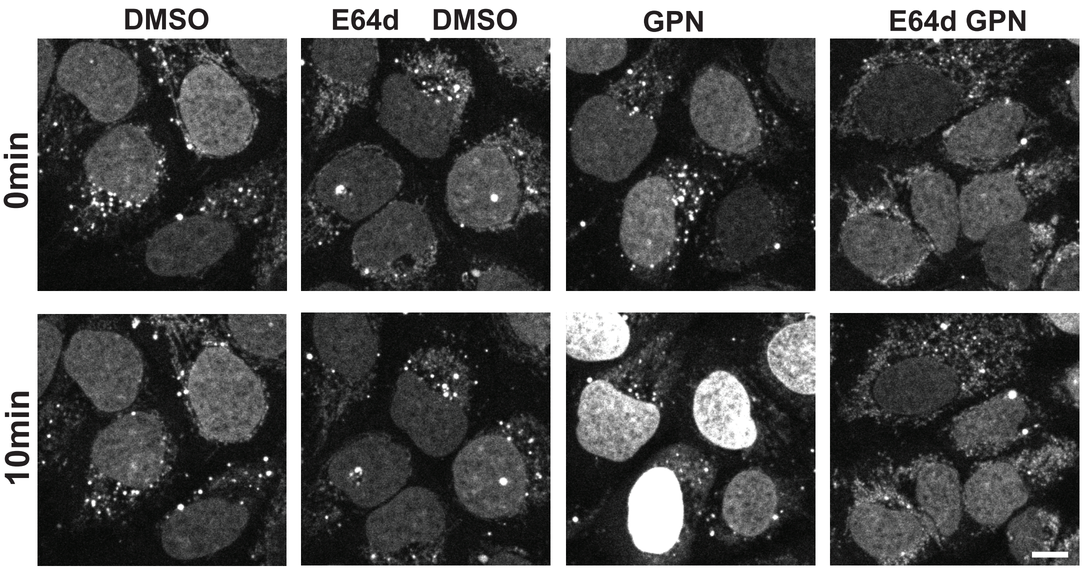


**Figure S1. Representative images from experiments in Figure 1D.** Images of SYTOX fluorescence in U2OS cells before and 10min after indicated treatment are shown. Scale bars, 10µm.

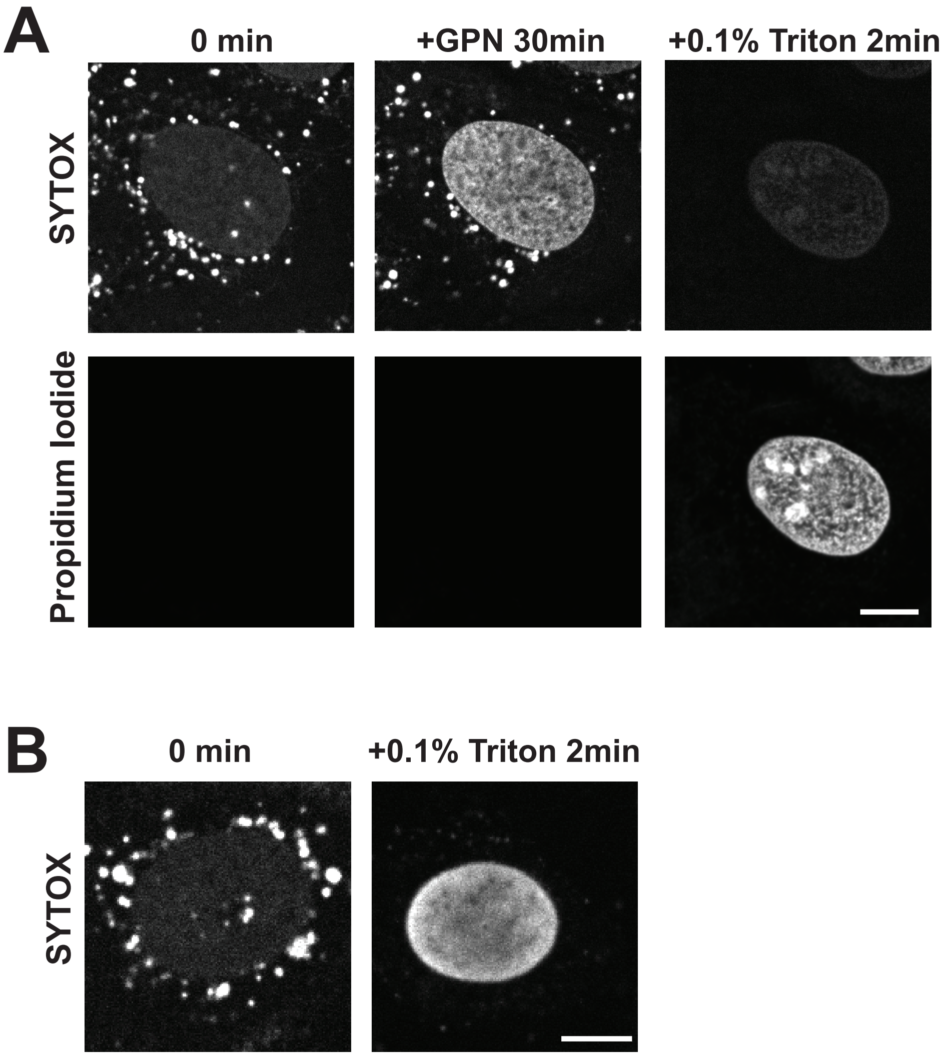


**Figure S2. Plasma membrane permeability is not compromised by GPN.** (A) U2OS cells were loaded with SYTOX Green followed by 4 hr chase as described. Propidium Iodide was added in the imaging media before imaging. Cells were treated with 0.2mM GPN for 30 min and then 0.1% Triton was added. Propidium Iodide intensity increased after addition of Triton and SYTOX was quenched by Propidium Iodide. (B) U2OS cells were loaded with SYTOX Green. 0.1% Triton was added to the cells in the absence of Propidium Iodide and nuclear SYTOX intensity increased. Scale bars, 10μm.

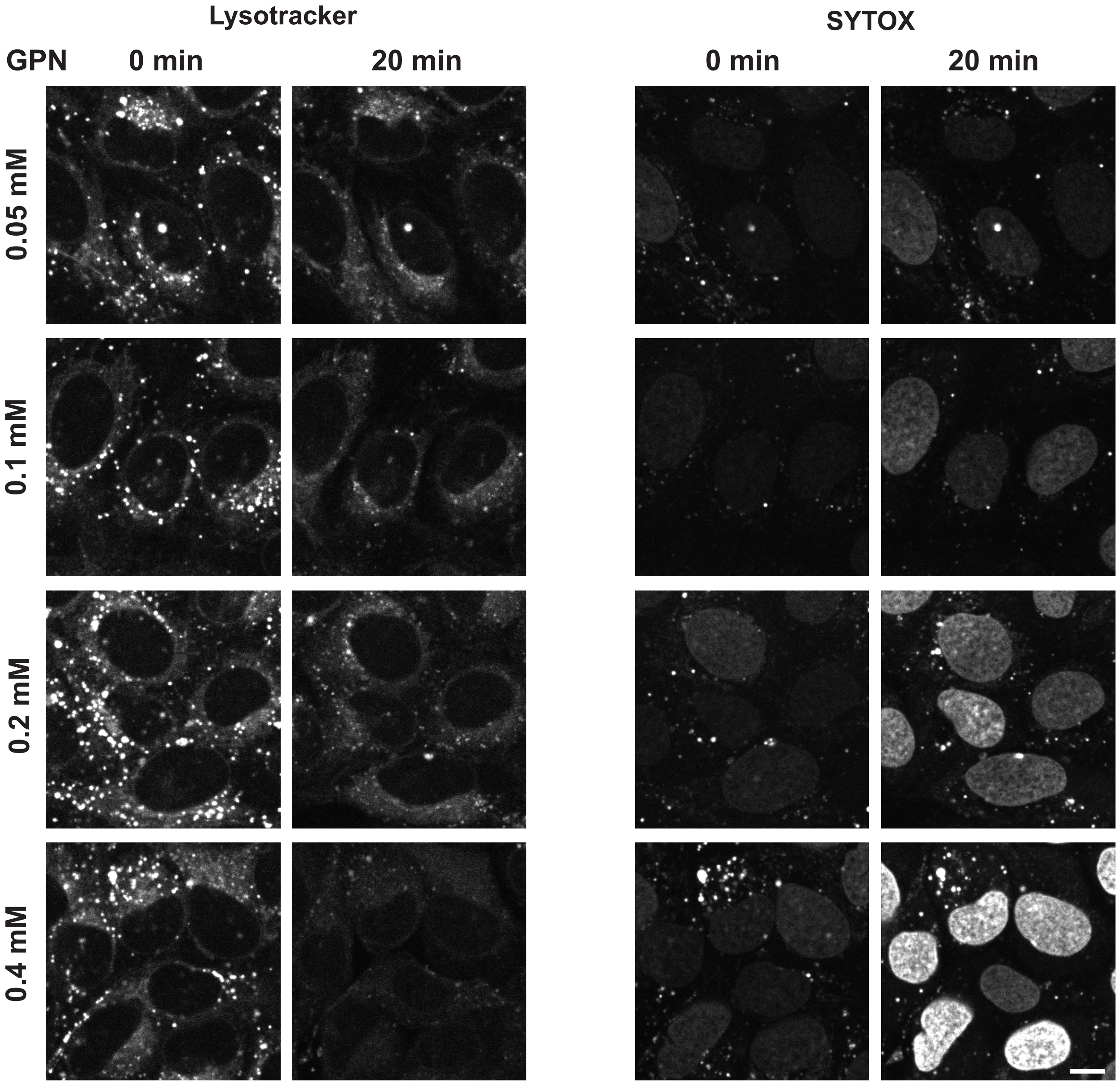


**Figure S3. Representative images from experiments in Figure 1F and 1G.** Images of SYTOX fluorescence in U2OS cells before and 20 min after GPN at indicated concentrations are shown. Scale bars, 10μm.

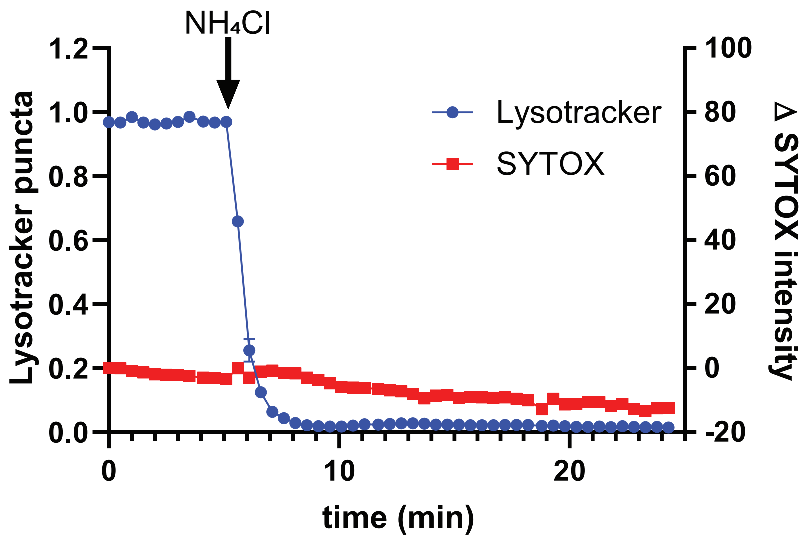


**Figure S4. Effects of $NH_4Cl$ on Lysotracker and SYTOX release.** U2OS cells were loaded with SYTOX Green and stained with Lysotracker before imaging. 20mM $NH_4Cl$ was added to cells at indicated time point.

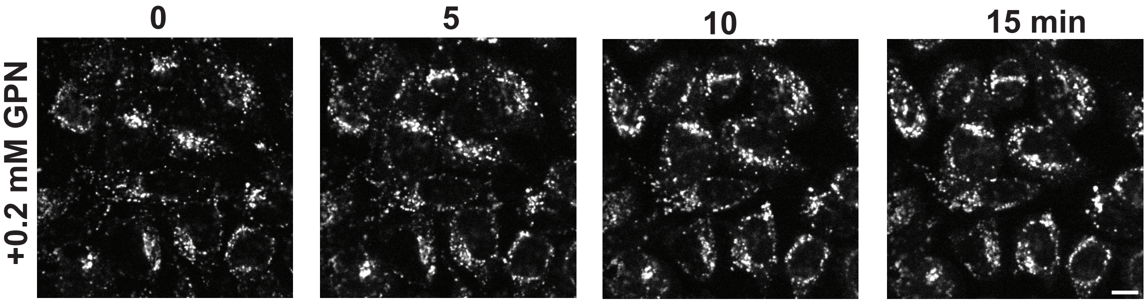


**Figure S5. 0.2mM GPN does not trigger loss of lysosomal dextran.** Time course images of U2OS cells loaded with 10kD cascade blue dextran treated with GPN for indicated time points. Scale bars, 10μm.

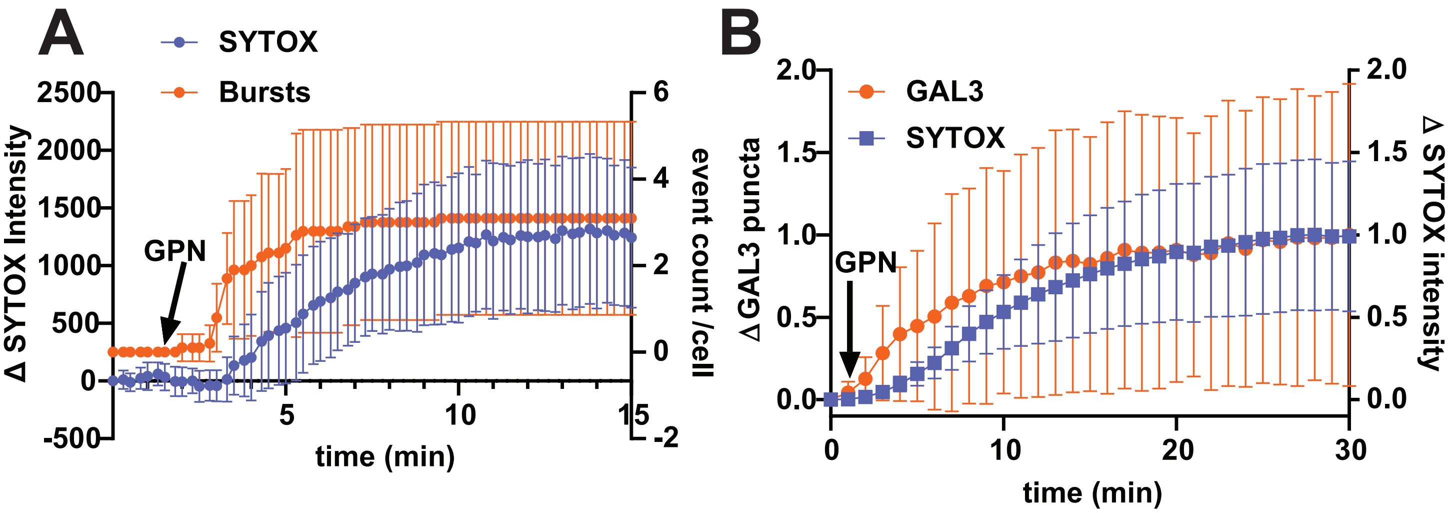


**Figure S6. Lysosomal bursts precede and correlate with nuclear SYTOX accumulation.** U2OS cells loaded with SYTOX Green and imaged live at 30s interval during exposure to GPN. Cumulative bursts (mean ± SD) from 10 cells are plotted with intensity change (mean ± SD, arbitrary unit) of nuclear SYTOX in (A). (B) U2OS cells expressing mCherry-GAL3 were loaded with SYTOX Green and imaged live at 30s interval during exposure to GPN. Normalized GAL3 puncta number (mean ± SD) from 10 cells are plotted with normalized intensity change (mean ± SD) of nuclear SYTOX in (B).

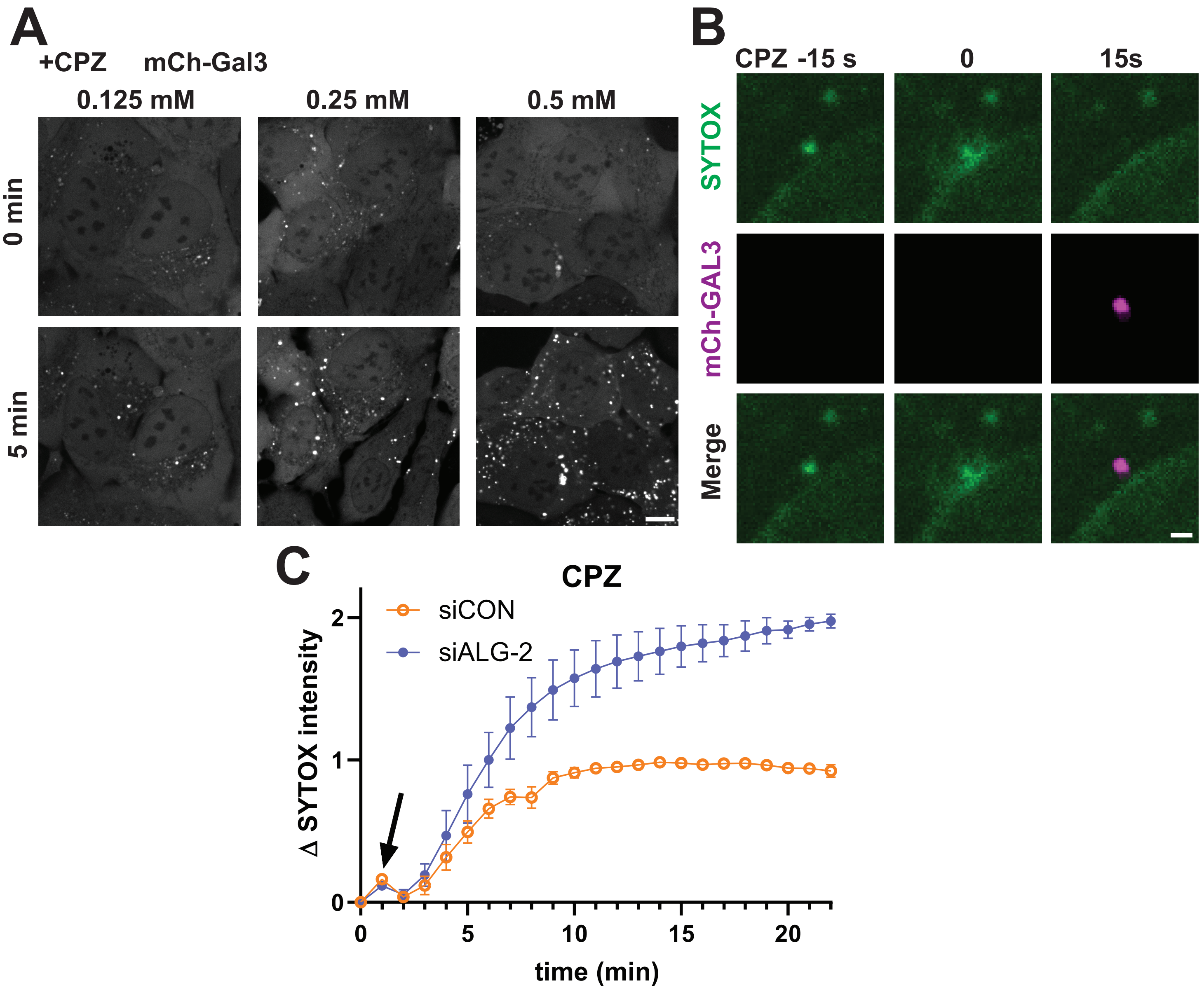


**Figure S7. ALG-2 deficient cells are more sensitive to CPZ-induced lysosomal damage.** (A) U2OS cells expressing mCherry-Gal3 were imaged live at 1min interval after exposure to CPZ at indicated concentrations. Representative images before and 5 min after CPZ. (B) U2OS cells expressing mCherry-Gal3 were loaded with SYTOX and imaged live at 15s interval during exposure to CPZ. Shown is representative burst of SYTOX release coinciding with abrupt Gal3 accumulation. Burst was set at time 0. (C) HeLa cells transfected with siCON or siALG-2 were loaded with SYTOX and imaged live at 1 min intervals before and after adding 0.25mM CPZ. Changes in nuclear SYTOX intensity (mean ± SEM, normalized to the mean value of plateau nuclear SYTOX intensity change in siCON group; n>200 cells for each condition from 3 independent experiments) were measured and plotted for each time point. Scale bars, 10 μm (A) and 1 μm (B).

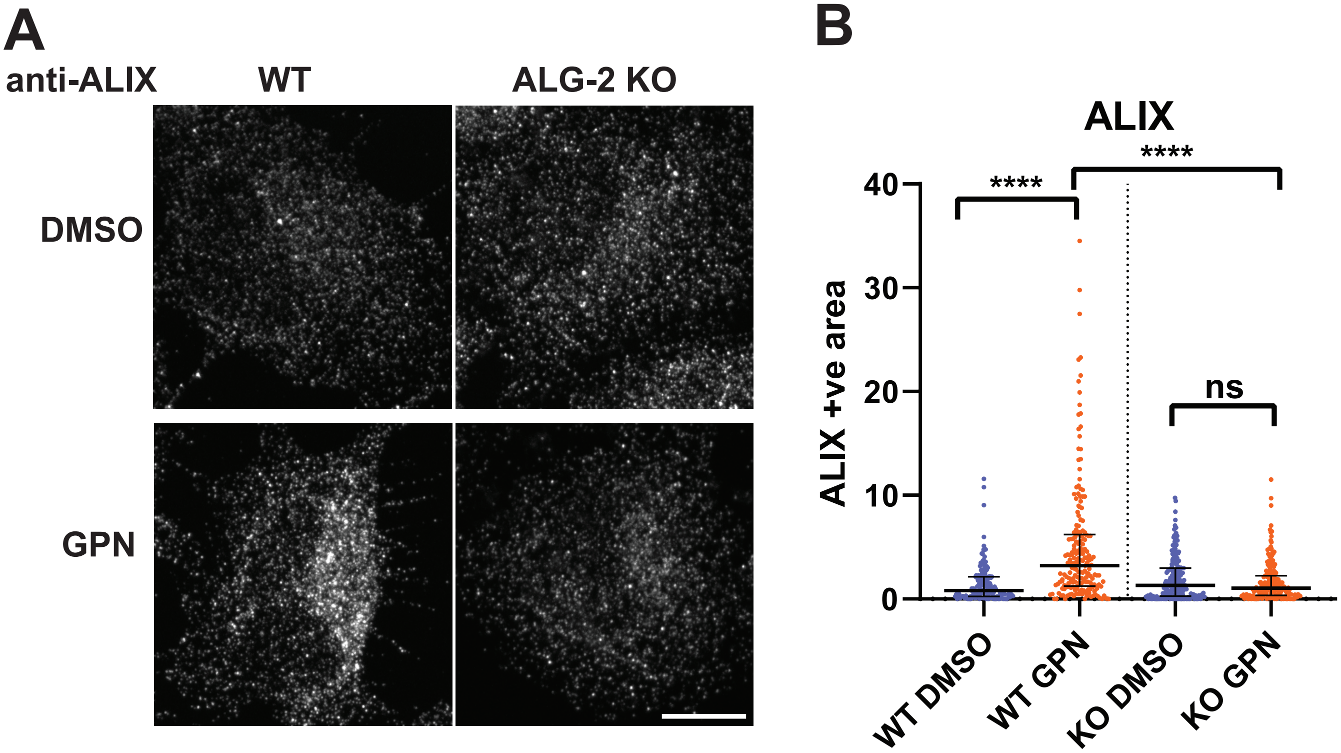


**Figure S8. GPN-triggered recruitment of ALIX to lysosomes depends on ALG-2.** (A) WT and ALG-2 KO HeLa cells were treated with DMSO or GPN for 10min and stained for ALIX. (B) ALIX positive area. Median with interquartile range (more than 150 cells for each condition from 3 independent experiments) of puncta area per cell are plotted. **** denotes P<0.0001; ns denotes P>0.9999 by Kruskal–Wallis test. Scale bars, 10μm.

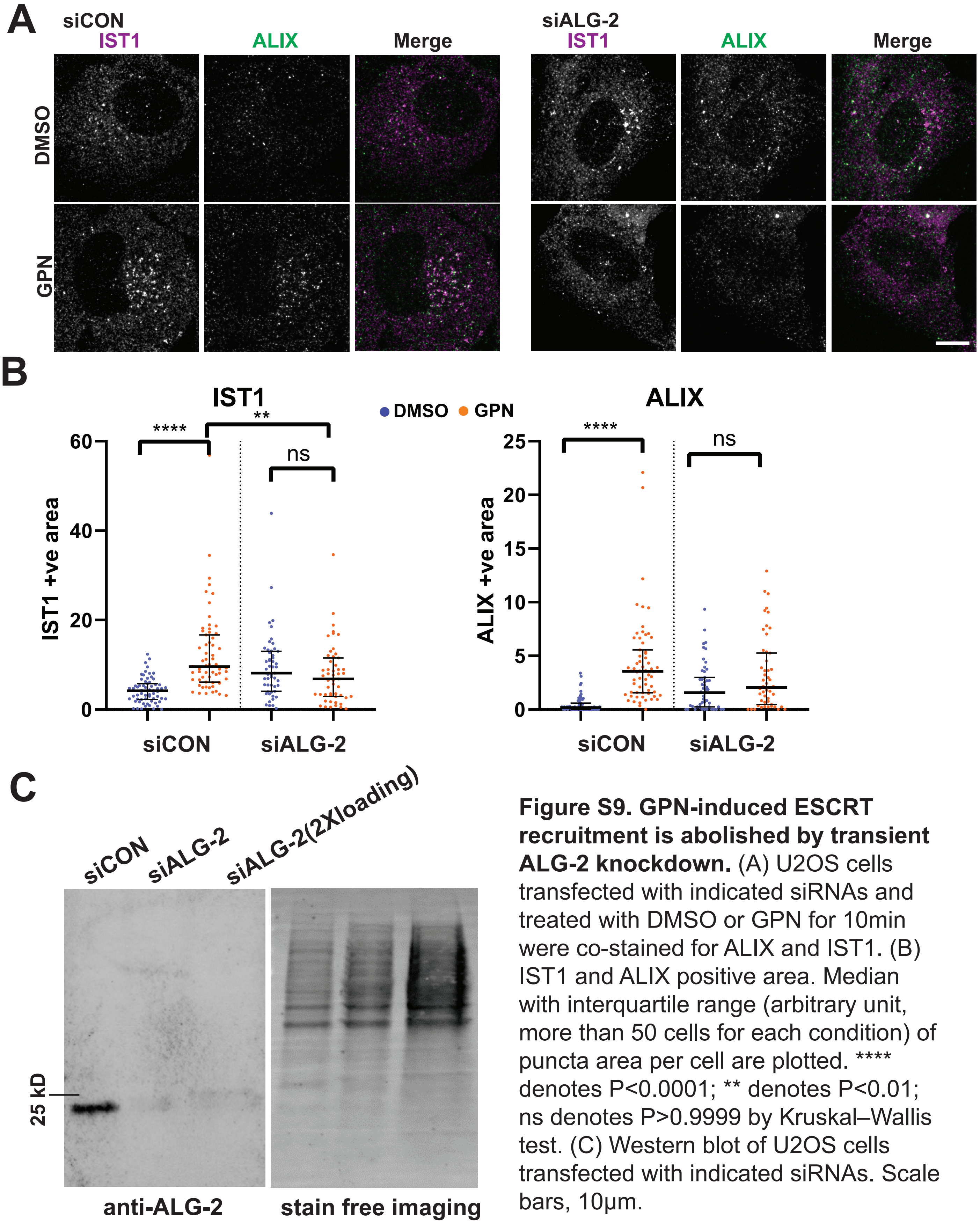


**Figure S9. GPN-induced ESCRT recruitment is abolished by transient ALG-2 knockdown.** (A) U2OS cells transfected with indicated siRNAs and treated with DMSO or GPN for 10min were co-stained for ALIX and IST1. (B) IST1 and ALIX positive area. Median with interquartile range (arbitrary unit, more than 50 cells for each condition) of puncta area per cell are plotted. **** denotes P<0.0001; ** denotes P<0.01; ns denotes P>0.9999 by Kruskal–Wallis test. (C) Western blot of U2OS cells transfected with indicated siRNAs. Scale bars, 10μm.

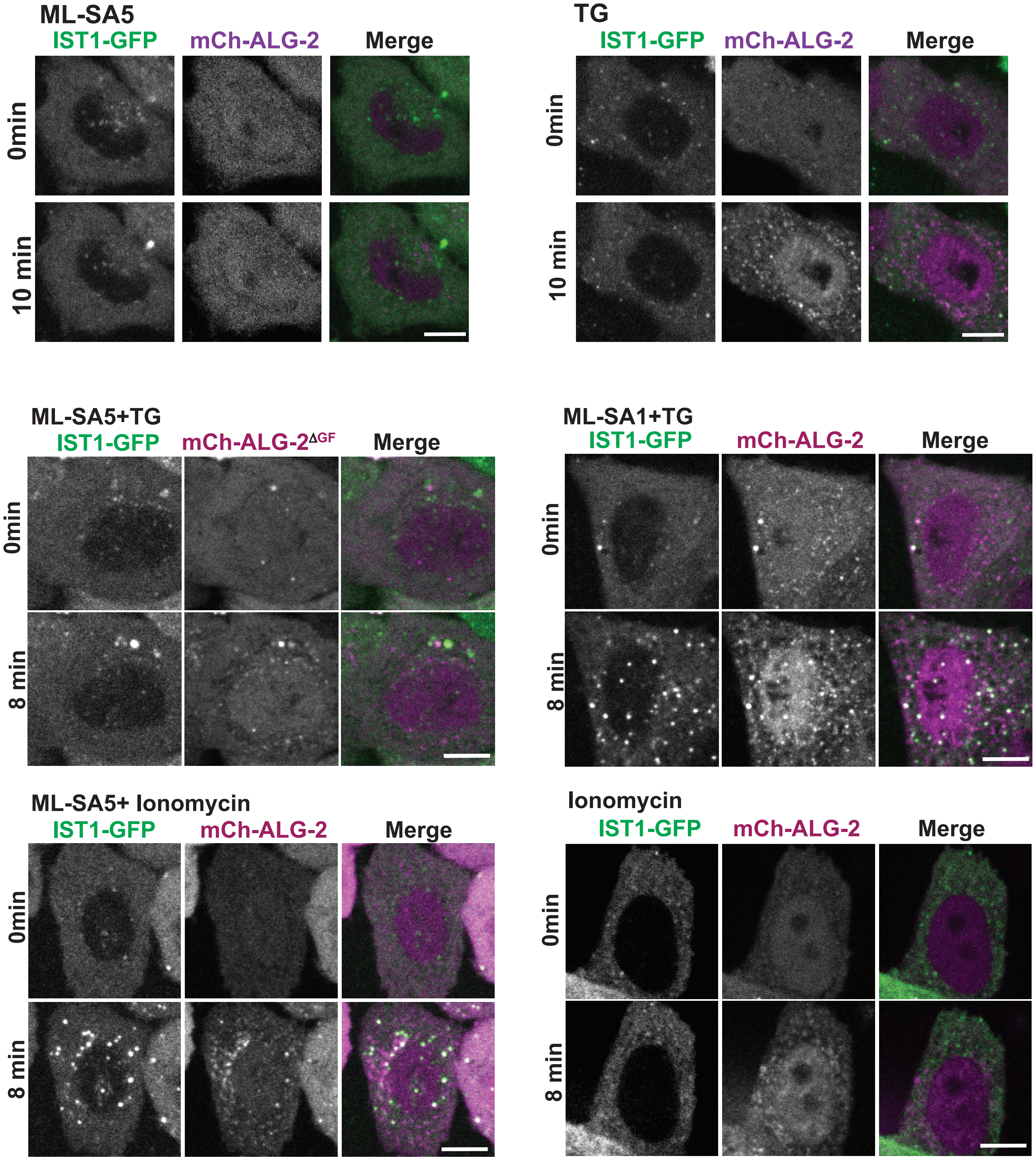


**Figure S10. Extended information related to Fig. 5.** ALG-2 KO HeLa cells co-transfected with IST1-GFP and mcherry-ALG-2 or mcherry-ALG-2$^{\Delta GF}$ were imaged live before and after: treatment with ML-SA5, TG, co-treatment with ML-SA5 and TG; co-treatment with ML-SA1 and TG; ML-SA5 and Ionomycin; or treatment with Ionomycin as lablled. Scale bars, 10μm.

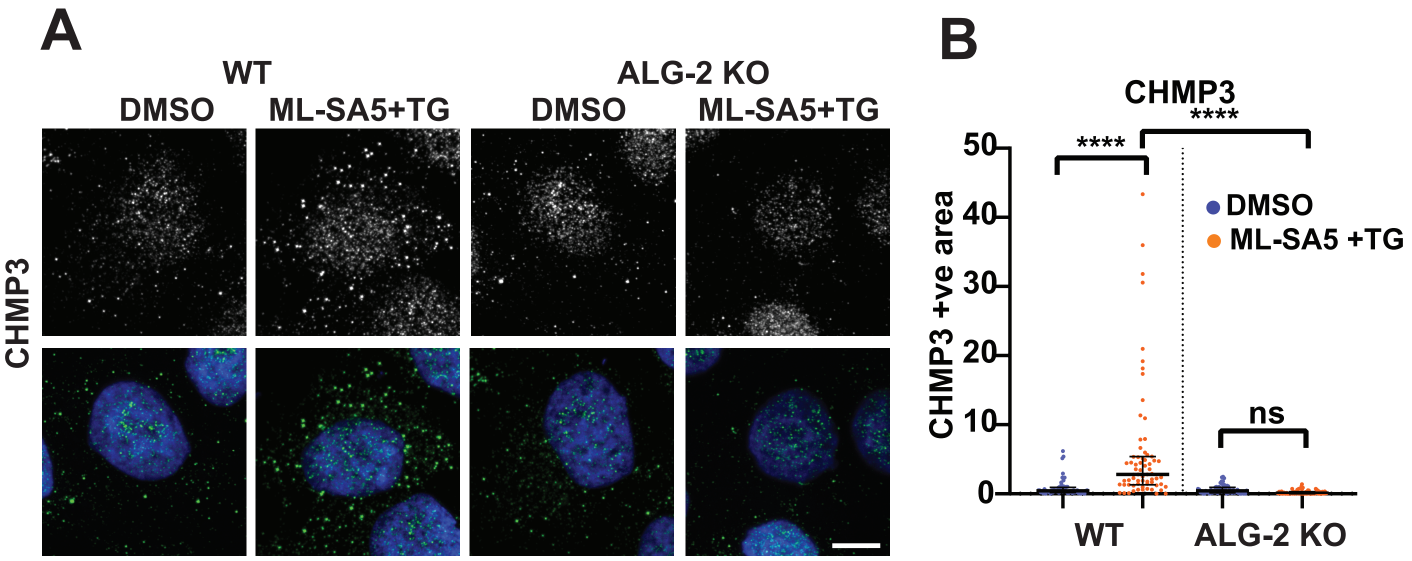


**Figure S11. Juxtalysosomal $Ca^{2+}$ triggered CHMP3 recruitment to lysosomes is ALG-2 dependent.** (A) WT and ALG-2 KO HeLa cells treated with DMSO or ML-SA5 with TG for 10min were stained for CHMP3. DAPI staining is also shown in bottom panel. (B) Cytosolic CHMP3 positive area. Median with interquartile range (more than 50 cells for each condition) of puncta area per cell are plotted. **** denotes P<0.0001; ns denotes P>0.9999 by Kruskal–Wallis test. Scale bars, 10μm.

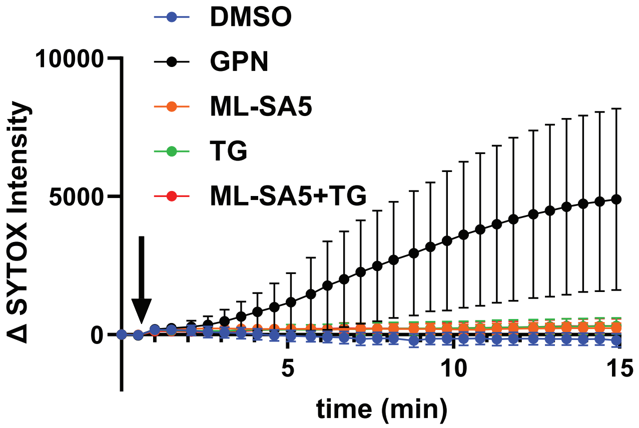


**Figure S12. ML-SA5 and TG do not cause SYTOX leak.** U2OS cells loaded with SYTOX Green were imaged live at 30s interval during indicated treatments. Changes in nuclear SYTOX Green intensities (arbitrary unit; mean ± SD; n>90 cells for each condition) were measured and plotted for each time point.

Movie 1
A whole field timelapse recording that corresponds to the time series shown in Fig. 1B. GPN was added to U2OS cell with SYTOX-loaded lysosomes at indicated time point. Images were captured every 15 s.

Movie 2
Timelapse recording that corresponds to the time series shown in Fig. 4B. ALG-2 KO HeLa cells were transfected with mCherry-ALG-2 or mCherry-ALG2$^{E47A/E114A}$ and loaded with Oregon Green 70 kD dextran to label lysosomes. GPN was added to cells at indicated time point. Images were captured every 1 min.

Movie 3
Timelapse recording that corresponds to the time series shown in Fig. 4C. ALG-2 KO HeLa cells were co-transfected with IST1-GFP and mcherry-ALG-2(FL) or ΔGF122. GPN was added to cells at indicated time point. Images were captured every 15 s.
